# Evolutionary genomics predicts adaptive genetic and plastic gene expression responses to climate change in a key alpine forest tree species

**DOI:** 10.1101/2023.07.11.548483

**Authors:** Zhiqin Long, Yupeng Sang, Jiajun Feng, Xinxin Zhang, Tingting Shi, Lushui Zhang, Kangshan Mao, Loren H. Rieseberg, Jianquan Liu, Jing Wang

## Abstract

Despite widespread biodiversity loss, our understanding of how species and populations will respond to accelerated climate change remains limited. In this study, we predict the evolutionary responses of *Populus lasiocarpa*, a key alpine forest tree species primarily found in the mountainous regions of a global biodiversity hotspot, to climate change. We accomplish this by generating and integrating a new reference genome for *P. lasiocarpa*, re-sequencing data for 200 samples, and gene expression profiles for leaf and root tissue following exposure to heat and waterlogging. Analyses of the re-sequencing data indicate that demographic dynamics, divergent selection, and long-term balancing selection have shaped and maintained genetic variation within and between populations over historical timescales. In examining genomic signatures of contemporary climate adaptation, we found that haplotype blocks, characterized by inversion polymorphisms that suppress recombination, play a crucial role in clustering environmentally adaptive variations. Comparison of evolved and plastic gene expression show that genes with expression plasticity generally align with evolved responses, highlighting the adaptive role of plasticity. Lastly, we incorporated local adaptation, migration, genetic load, and plasticity responses into our predictions of population-level climate change risks. Our findings reveal that western populations, primarily distributed in the Hengduan Mountains—a region known for its environmental heterogeneity and significant biodiversity—are the most vulnerable to climate change and should be prioritized for conservation and management. Overall, our study advances understanding of the relative roles of long-term natural selection, local environmental adaptation, and immediate plasticity responses in driving evolutionary adaptation to climate change in keystone species.

## INTRODUCTION

Environmental changes driven by ongoing anthropogenic climate change have led to shifts in species ranges, population decline, and even extinction, establishing it as a significant driver of global biodiversity loss. In this context, a critical priority for biodiversity conservation is understanding and monitoring how keystone species in the local ecosystem respond to rapid climate change [1]. Genomic data serves as one of the primary repositories for recording genetic adaptation to past environmental fluctuations and offers a potential useful tool for predicting the evolutionary potential for future adaptation in the context of climate change [2]. Integration of ecological and climate genomics is needed to better comprehend and forecast the impacts of climate change on diversity at the genome, population, and species levels [3].

In the face of unprecedented environmental changes, natural populations-particularly plants with limited mobility-can persist *in situ* through genetically based evolutionary changes and/or environmentally induced phenotypic and gene expression plasticity. Alternatively, they may face the risk of extirpation. In the context of genetically adaptive evolution, various evolutionary processes, including mutation, natural selection, and recombination, play pivotal roles in shaping responses to climate change [4]. The ability to rapidly respond to environmental changes largely depends on the amount of standing adaptive genetic variation, including adaptive single nucleotide polymorphisms (SNPs), short insertions/deletions (indels), and large structural variants (SVs). These variants are largely maintained by long-term balancing selection and/or divergent selection within and among populations [5]. Notably, one type of SV, inversion polymorphisms, has been repeatedly linked to local ecological adaptation in species such as mimetic butterflies [6], monkey flowers [7], sunflowers [8], and deer mouse [9]. Inversions suppress recombination in heterozygotes, helping to keep cassettes of locally adapted alleles together in the face of maladaptive gene flow.

In addition, gene expression variation is highly sensitive to environmental effects, and thus environmentally induced expression plasticity is hypothesized to be crucial for facilitating an immediate response to rapid environmental change [10]. However, the relationship between evolutionary adaptation and environmental plasticity remains poorly understood. For instance, plastic changes can be adaptive if the direction of plastic response aligns with the direction of natural selection, serving as a steppingstone to reach a fitness optimum in the new environment. Conversely, plasticity is maladaptive if the direction of plastic responses is opposite to the direction of evolved response, potentially moving individuals further away from the fitness optimum [10–13]. Therefore, elucidating the relative roles of evolutionary genetic adaptation, immediate plastic expression changes, and their interactive roles for organismal adaptation to new environments is crucial for a comprehensive understanding of the mechanisms driving adaptation, vulnerability, and resilience during the responses of natural populations to future climate change.

Forest ecosystems serve as habitats for biodiversity and provide environmentally friendly products for humans. Functioning as architects of ecosystems, trees efficiently act as carbon sinks, playing a crucial role in the global carbon cycle [14]. However, due to their lengthy lifespans, most trees face significant challenges in keeping up with rapid climate change, particularly considering that the threat is likely to manifest within the lifetime of a single individual [15, 16]. The mountains of Southeast Asia, with their highly complex and heterogeneous environments, are recognized as global biodiversity hotspots, notable for exceptional species endemism [17]. Given the increasing environmental fluctuations, species in this region are particularly vulnerable to maladaptation under future climate change risks [18]. *Populus lasiocarpa*, a dominant and endemic dioecious tree species, is distinguished by its large leaves within the genus *Populus* and is primarily distributed in mountain regions surrounding the Sichuan Basin in Southeast Asia [19]. Natural populations are found in the Hengduan Mountains (west), Qinling Mountains (north), Daba and Wushan Mountains (east), and Yunnan-Guizhou Plateau (south) (Fig. 2A). This ring-shaped distribution makes *P. lasiocarpa* an ideal system for exploring the evolutionary forces and genetic mechanisms underlying climate adaptation and stress tolerance in natural forest populations in these mountainous regions [20, 21]. There is a pressing need for comprehensive prediction and assessment of a species’ population-level vulnerability to future climate change, particularly for keystone forest trees endemic to this area.

In this study, we constructed a *de-novo* chromosome-level reference genome of *P. lasiocarpa* and subsequently acquired whole-genome re-sequencing data from 200 individuals representing 22 populations across its natural distribution range (Fig. 2A; Supplementary Fig. 1). Additionally, we obtained RNA-seq data for experimental seedlings from western and eastern populations, reflecting adaptation to distinct thermal and waterlogging environments under both control and stress conditions in leaf and root tissues. With this comprehensive genomic and transcriptomic dataset, our objectives are as follows: (1) investigate how demographic histories, geography, and natural selection have influenced the patterns of genomic variation; (2) dissect the genetic basis and identify potential adaptive variants facilitating local environmental adaptation; (3) examine the relative roles and intricate relationships between genetically-based evolutionary changes and environmentally-induced expression plasticity underpinning responses to changing environments; (4) predict the future vulnerability of populations to climate change, and identify those populations at the highest risk and that require urgent management actions.

## RESULTS

### High-quality reference genome assembly of *P. lasiocarpa*

A multifaceted sequencing approach, including 49.95 Gb of Nanopore sequencing reads (∼119x), 36.37 Gb of Illumina reads (∼87x), and 62.52 Gb of high-throughput chromosome conformation capture (Hi-C) reads (∼149x), was used for chromosome-scale genome assembly of *P. lasiocarpa* (Supplementary Tables 1). After initial genome assembly, multiple rounds of polishing and redundant sequences filtering (Supplementary Table 2), the final assembly captured 419.54 Mb of genome sequences, with a contig N50 size of 9.19 Mb and 99.23% of the assembled sequences anchored to 19 pseudo-chromosomes (Supplementary Fig. 2; Supplementary Table 3). The completeness and accuracy of the assembled genome were validated and supported by a high proportion (97.9% and 97.8%, respectively) of recalled single-copy orthologs from the Benchmarking Universal Single-Copy Orthologs (BUSCO)[22] and Illumina short reads mapping rate (Supplementary Tables 4,5).

We subsequently annotated repetitive elements, protein-coding genes and non-coding RNAs. In total, transposable elements (TEs) made up 40.2% of the genome, with retrotransposons and DNA transposons representing 20.8% and 15.9% of the genome, respectively (Supplementary Table 6). After using a comprehensive strategy combining homology-, transcriptome- and *ab initio*-based prediction, a total of 39,008 protein-coding genes were annotated with high confidence (Figure 1; Supplementary Tables 7,8). Among the predicted protein-coding genes, 93.96% could be annotated through at least one of the public protein-related databases (Pfam, NR, InterPro, KEGG, Swiss-Prot, KOG, COG, TrEMBL and GO) (Supplementary Table 9). In addition, we predicted 5,322 non-coding RNAs containing ribosomal RNAs, transfer RNAs, microRNAs, and small nuclear RNAs (Supplementary Table 10).

**Fig. 1.**
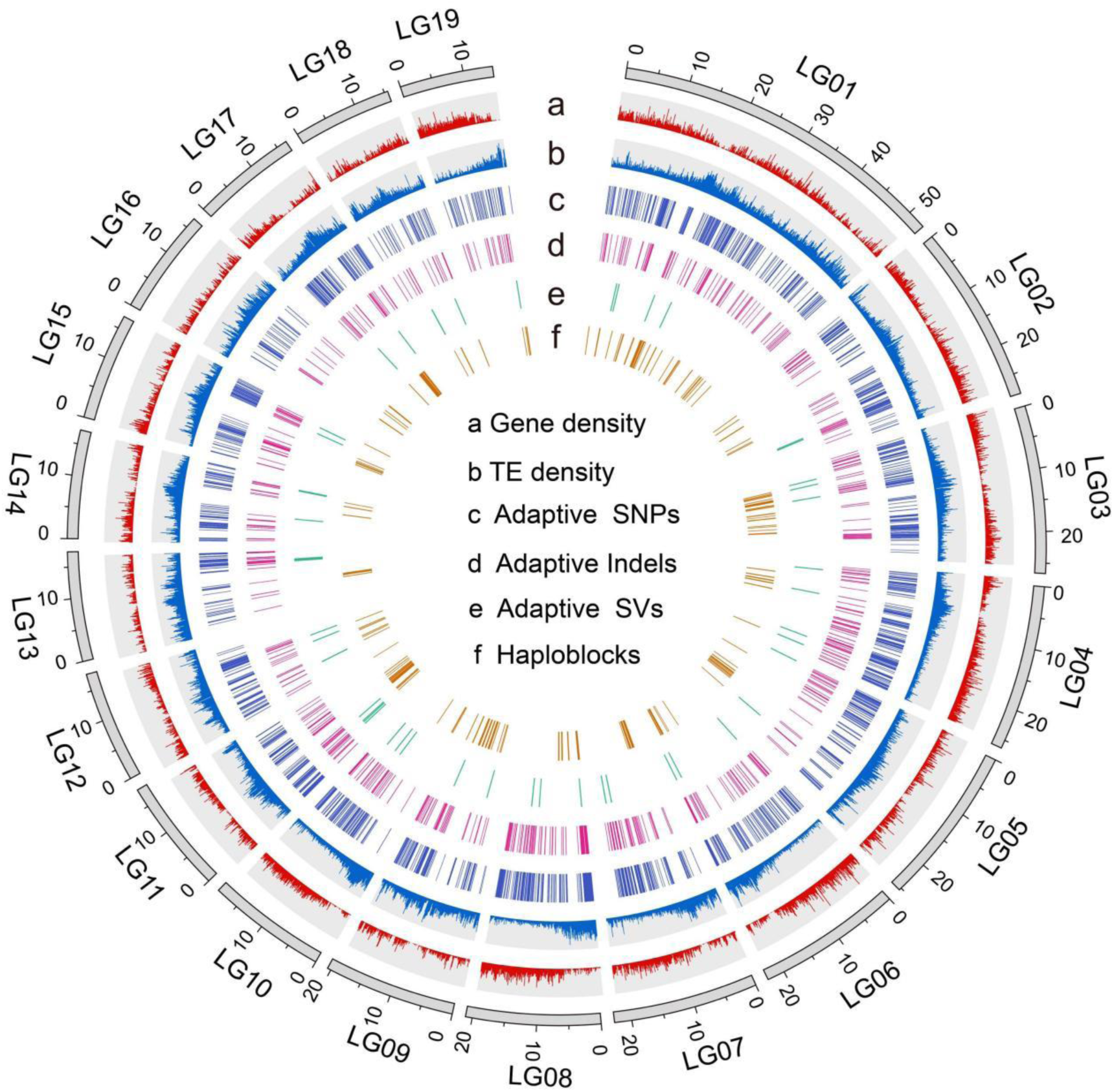
Genomic features of *P. lasiocarpa* reference genome. The Circos plot displays gene density (**a**), transposable element (TE) density (**b**), distribution of putatively climatic-adaptive SNPs (**c**), INDELs (**d**), SVs (**e**), and the distribution of haploblocks (**f**).

### Population genetic analyses offer insights into evolutionary history

To examine the relationships and divergence among populations of *P. lasiocarpa,* we conducted re-sequencing of 200 accessions from 22 populations spanning its natural distribution in Southwest China (Fig. 2A1). The sequencing was performed at an average depth of approximately 25×, with a mapping coverage of 95% (Supplementary Table 11). Following a series of rigorous filtering steps (see Methods), we identified a total of 10,567,146 high-quality SNPs, 1,534,259 indels, and 64,954 SVs. To validate the quality of SVs detected in this study, we inferred population structure using a randomly sampled set of 156,324 linkage disequilibrium (LD)-pruned SNPs and 9,989 SVs. Both datasets exhibited highly consistent results (Fig. 2A2; Supplementary Fig. 3).

**Fig. 2.**
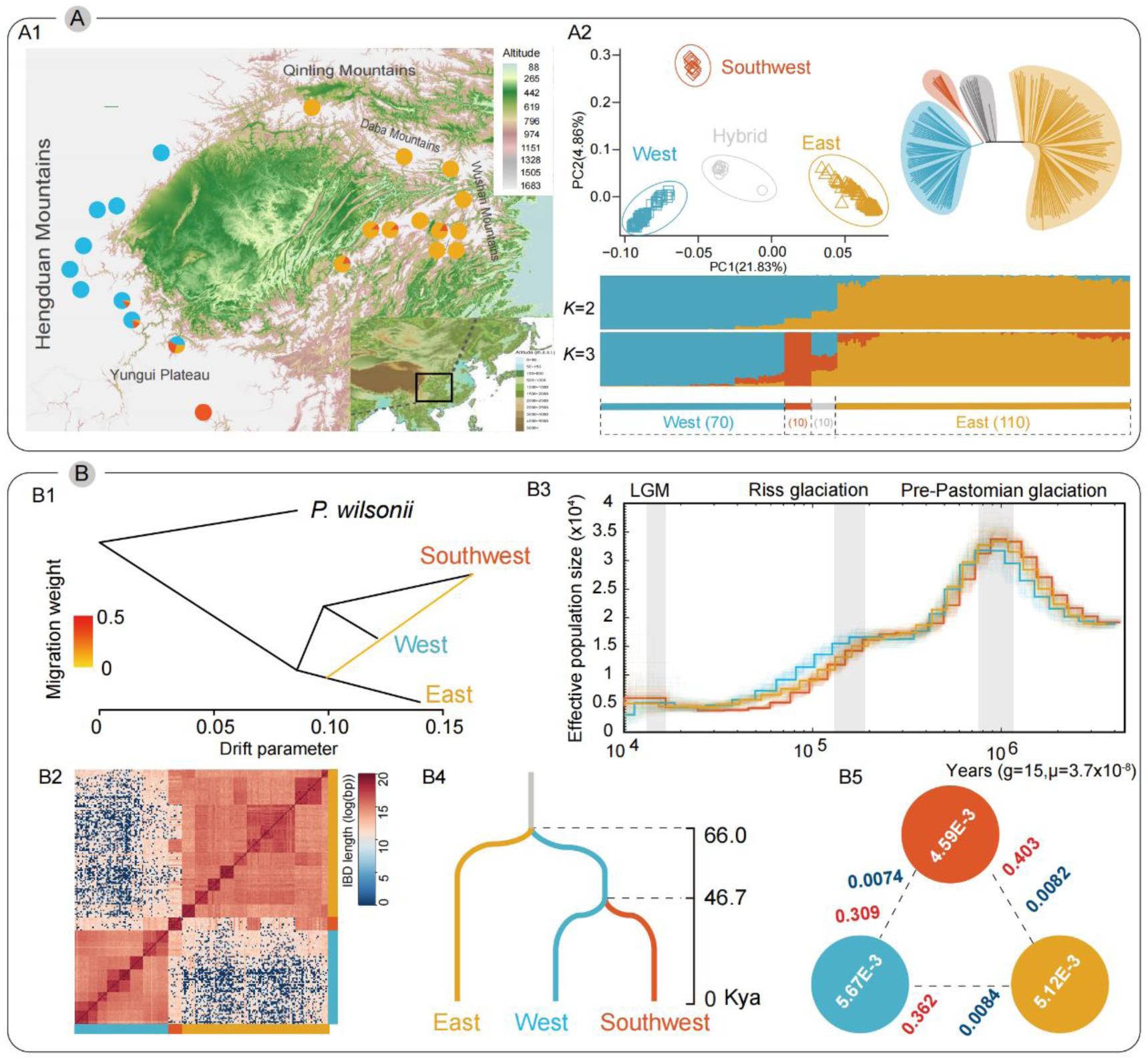
Population genomic analyses of *P. lasiocarpa*. **(A1)** Geographic distribution of sampled *P. lasiocarpa* populations, with colors representing ancestral components inferred by ADMIXTURE at *K*=3. **(A2)** Neighbor-joining (NJ) phylogenetic tree, PCA, and genetic cluster plots for a *K*=2 and *K*=3, where the height of each colored segment represents the proportion of the individual’s genome derived from inferred ancestral lineages. The number of individuals in each group is shown within the dashed-line box. **(B1)** Population splits and migrations among three groups of *P. lasiocarpa* using TreeMix analysis. **(B2)** Estimated haplotype sharing among individuals of *P. lasiocarpa*. The heatmap color reflects the total length of identity-by-descent (IBD) among individuals. **(B3)** Demographic history revealed by the PSMC method. For each lineage, the median estimate for all 10 selected individuals is shown with a bold line, while 100 bootstrap replicates (ten replicates for each of the 10 samples) are shown with faint lines. The gray vertical bars indicate the periods of the Last Glacial Maximum, Riss Glaciation, and Pre-Pastonian Glaciation. **(B4)** Divergence times of three groups, with timelines of splits inferred by SMC++. **(B5)** Estimation of nucleotide diversity (π), relative genetic divergence (*F*_ST_), and absolute genetic divergence (*d*_XY_) within and between the three lineages (red: Southwest; blue: West; orange: East).

Both neighbor-joining (NJ) phylogenetic tree and principal component analysis (PCA) unveiled the presence of four genetic clusters within *P. lasiocarpa* corresponding to the West, East, Southwest, and Hybrid groups (Fig. 2A2). The West group consists of 70 individuals from eight populations primarily distributed in the Hengduan Mountains in western areas of the Sichuan Basin. The East group encompasses 110 individuals from 12 populations distributed in the Daba and Wushan Mountains in eastern areas of the Sichuan Basin. In contrast to the two main groups of western and eastern populations, the Southwest and Hybrid groups only include 10 individuals separately. Additionally, clustering analysis conducted by ADMIXTURE identified the strongest support for *K*=3 populations, thereby supporting the separation of West, East, and Southwest groups of populations (Fig. 2A2). Due to the high level of inter-population admixture in the Hybrid group, we subsequently excluded this group from the following analysis.

To delve further into divergence and demographic histories, we initially employed TreeMix to infer the phylogenetic tree topology among the three main groups. The results indicated that the Southwest and West groups shared a recent common ancestor, with a migration edge suggesting gene flow from East to Southwest group (Fig. 2B1). Another haplotype sharing analysis revealed that individuals within the same group exhibited more and longer identity-by-descent blocks compared to those from different groups. Notably, there was a lower and shorter haplotype-sharing pattern observed between the West and East groups than between each of them and the Southwest group (Fig. 2B2).

Subsequently we applied the pairwise sequentially Markovian coalescent (PSMC) model to infer the fluctuations in effective population size (*N_e_*) over time in each group. The results supported a shared decline of populations among these groups (Fig. 2B3). Recognizing the limited power of recent *N_e_* trajectory inference for PSMC, we employed SMC++ to explore the more recent demographic history of these populations. Compared to the continuous reduction in *N_e_* observed in the Southwest group, both the West and East groups experienced recent population expansion following a shared bottleneck after the Last Glacial Maximum (Supplementary Fig. 4). We further estimated the divergence time between the population groups, revealing that the West group diverged from the East group approximately 66 thousand years ago (Kya). The subsequent divergence time between the Southwest and West group was estimated at approximately 47 Kya (Fig. 2B4).

Furthermore, we investigated and compared patterns and levels of nucleotide diversity (π) and genetic differentiation (*F_ST_* and *d_XY_*) among the populations. Consistent with the slower rate of LD decay observed in the Southwest group of populations (Supplementary Fig. 5), nucleotide diversity was also found to be lowest (π=4.59×10^-3^) compared to the West (π=5.67×10^-3^) and East (π=5.12×10^-3^) groups of populations (Fig. 2B5), aligning with the corresponding bottleneck observed in this group. Finally, after adjusting for the varying number of individuals in each group, we randomly selected 10 individuals from the three groups and obtained highly consistent results in the analysis of π, LD, *F_ST_*, *d_XY_* and recent demographic history (Supplementary Fig. 6).

### Genome-wide signatures of divergent and balancing selection

Ancient polymorphisms may be sustained through long-term balancing selection during population divergence, resulting in genomic regions with increased genetic diversity within and among nascent populations[23]. Alternatively, ancestral balanced polymorphisms might be sorted unequally across descendant lineages by selection and drift, leading to genomic regions with increased divergence [24, 25]. These evolutionary forces can give rise to genomic regions with heterogeneous patterns of relatedness, potentially confounding overall population structure. To unveil the evolutionary processes shaping the landscape of genomic divergence, we initially excluded the Southwest group, given its smaller individual size compared to the other two main groups. We then characterized local population structure by estimating multidimensional scaling (MDS) in non-overlapping windows of 250 SNPs along the genome using a local PCA approach [26]. To identify outlier windows, we selected the top and bottom 0.5% windows in the MDS1 (Fig. 3A). In comparison with the genomic background windows, the two extreme windows exhibited distinct phylogenetic clustering topologies and population-wide relationships (Fig. 3B), suggesting that these regions are likely under long-term balancing selection (the top 0.5% windows) and divergent selection (the bottom 0.5% windows) respectively.

**Fig. 3.**
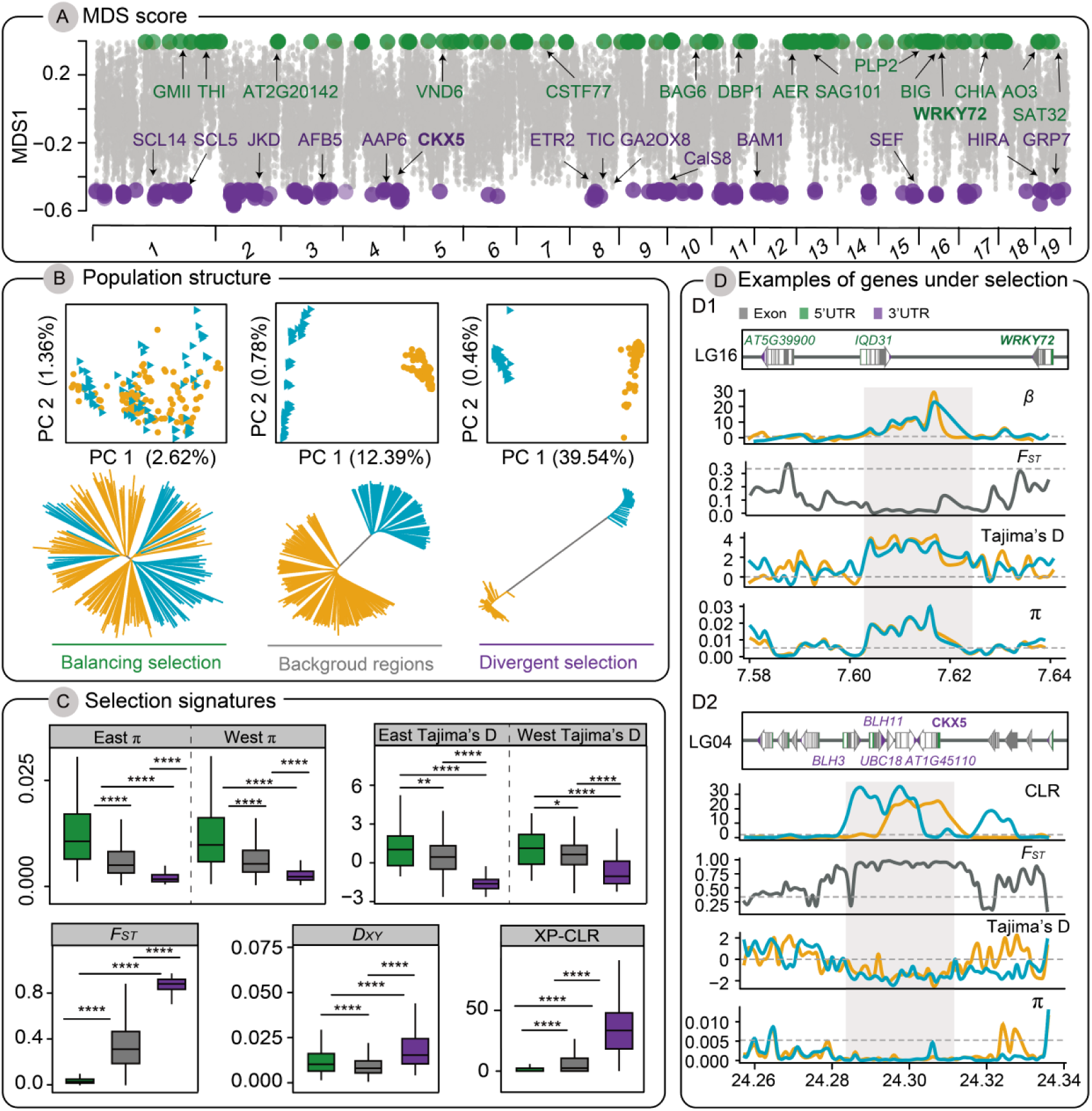
Identification of balancing and divergent selection regions. **(A)** Characterization of MDS (multidimensional scaling) outlier regions across the genome using local PCA. Each dot represents a window containing 250 SNPs. Candidate regions under balancing selection are colored green, while those under divergent selection are colored purple. **(B)** PCA and NJ tree analyses based on SNPs from balancing selection regions, background regions, and divergent selection regions. **(C)** Comparison of π, Tajima’s D, *F_ST_, d_XY_*, and XP-CLR among balancing selection regions (green), background regions (grey), and divergent selection regions (purple). Significance was determined using the Wilcoxon test (two-tailed), with asterisks indicating significance levels above the boxplot (\*\*\**P*<0.001, \*\**P*<0.01, \**P*<0.05, ns: *P*>0.05). **(D)** Distribution of β, CLR, *F*_ST_, Tajima’s D, and π around examples of candidate genes in balancing (D1) and divergent (D2) selection regions. Grey dashed lines represent the genome-wide significance level of the top 0.5% for each parameter. The structure of each candidate gene is shown in the upper box.

To further explore the potential evolutionary processes driving the divergent patterns of genomic variation in these regions, we compared the level of nucleotide diversity (π), Tajima’s D, both the relative (*F_ST_)* and absolute (*d_XY_*) genetic divergence, and the cross-population composite likelihood ratio statistics (XP-CLR) among the three types of windows. Consistent with the hypothesis of long-term balancing selection, we found that the top 0.5% windows showed significantly elevated π, more positive values of Tajima’s D, higher *d_XY_*, and lower *F_ST_* and XP-CLR values compared to genomic background windows. Consequently, we referred these windows as ‘Balancing selection (BLS) regions’ (185 BLS windows with a total length of 1.32 Mb; ∼0.32% of the assembled genome) (Fig. 3 and Supplementary Table 12). Further Gene Ontology (GO) functional enrichment analysis of the 166 candidate genes in BLS regions revealed categories significantly enriched in the metabolic process and biotic stress response, including thiamine, oxazoline and thiazoline metabolic and biosynthetic processes that play a crucial role in responding to DNA damage and pathogen attack in plants (Supplementary Fig. 7B). Specifically, multiple genes under balancing selection were orthologous to *Arabidopsis* genes associated with plant immune responses (Supplementary Fig. 8). For example, *Polas33741* (Fig. 3D1) was orthologous to *WRKY72*, which was found to regulate defense-related and jasmonic acid (JA) biosynthesis genes in many plants [27].

Conversely, we found that the bottom 0.5% windows exhibited strongly decreased π, more negative values of Tajima’s D, higher *d_XY_*, higher *F_ST_*, and XP-CLR compared to background windows and were henceforth referred to as ‘Divergent selection (DS)’ regions (185 DS windows with a total length of 1.76 Mb; ∼0.42% of the assembled genome) (Fig. 3 and Supplementary Table 13). GO enrichment analyses of 163 candidate genes in DS regions were significantly enriched in biological processes of xenobiotic stimulus and oxidative stress responses (Supplementary Fig. 7A). Specifically, many genes that showed high divergence and positive selection signatures were functionally related to organ development, rhythm control, and hypoxia adaptation (Supplementary Fig. 7C; Supplementary Fig. 9; Supplementary Table 13). For example, *Polas12612* (Fig. 3D2), the ortholog in *Arabidopsis* (*CKX5*) was found to be the downstream target gene of the phytochrome-interacting factor 4 (*PIF4*), which regulates photoperiod-dependent morphogenesis [28]; *Polas20885*, the ortholog (*TIC*) of which is involved in circadian rhythms control of plant [29]. Therefore, these candidate genes within the divergent selection regions likely underlie divergent ecological adaptation to the disparate mountain environments between the West and East lineages of *P. lasiocarpa*.

### The genomic signatures of environmental adaptation

To identify genomic variants associated with various environmental variables, we combined the Latent Factor Mixed Model (LFMM) [30] and Genome-wide Efficient Mixed Model Association (GEMMA) [31] to test the association between variant genotypes and each of the 29 environmental variables, while accounting for the confounding effects of population structure. We identify a total of 19,955 putatively adaptive variants from the overlapping results of LFMM and GEMMA (Supplementary Table 14), which are distributed widely across the genome (Supplementary Fig. 10). These variants include not only SNPs and short Indels (Supplementary Fig. 11) but also SVs, which are often overlooked (Supplementary Fig. 12). A functional enrichment analysis of the 2,044 genes with variants associated with environmental adaptation revealed significant enrichment in GO categories related to catabolic and metabolic processes (Supplementary Table 15), which are crucial for the survival of plants facing environmental changes [32, 33].

To further assess the robustness of the locally adaptive variants identified, we initially employed Mantel and partial Mantel tests to evaluate patterns of isolation-by-distance (IBD) and isolation-by-environment (IBE) for the environmental-associated “adaptive” variants and randomly selected “neutral” variants. Our results revealed that both adaptive and neutral variants exhibited strong patterns of IBD. However, after accounting for geographic effects with a partial Mantel test, we observed a significant correlation between pairwise genetic distance between populations (*F_ST_*/1-*F_ST_*) and environmental distance for adaptive variants. This was in contrast to the weak pattern of IBE observed in neutral variants (Supplementary Fig. 13). This suggest that the adaptive variants are primarily influenced by environmental gradients rather than neutral factors like population structure and geographic distance. Additionally, we employed a machine learning algorithms, gradient forest, to compare the shape of allele frequency turnover along the environmental gradient for adaptive and neutral variants from random sampling. Our findings indicated that the cumulative importance of allele frequency changes in the adaptive variants differed in shape and magnitude from those in the neutral reference datasets, suggesting that environmental variation more effectively explains allele frequency changes in the local adaptive variants compared to the reference variants (Supplementary Fig. 11D; Supplementary Fig. 12E).

### Haploblocks facilitates local adaptation

Large haploblocks, especially inversions, have been implicated to play a key role in local adaptation, primarily by facilitating the capture and linkage of multiple locally adaptive alleles through the suppression of recombination within co-inherited haplotype blocks [34–36]. Recent advancements in population genomic approaches have proven to be powerful and effective in detecting potential inverted regions across a diverse range of species [26]. In this study, we used an unbiased genome-wide scan approach with local principal component analyses (PCA) to characterize population structure and identify outlier regions using windows of 60 SNPs in size throughout the genome in *P. lasiocarpa.* To pinpoint putative inverted regions, we focused on genomic regions where individuals were distinctly clustered into three groups corresponding to the three possible inversion genotypes on the first principal component. The center cluster, showing the highest heterozygosity, was consistent with inversion heterozygotes (see Methods). We identified a total of 218 haploblocks, ranging in size from 10 to 650 kbp and encompassing 647 genes. GO categories within these haploblocks were notably enriched in ethylene-activated and phosphorelay signal transduction and signaling pathways (Supplementary Table 16).

To further substantiate the association of these haploblocks with structural variations, particularly inversions, we analyzed population-scale recombination rates, deleterious mutation loads (the ratio of zero-fold to four-fold variants), genetic divergence between populations (*F_ST_*), and overall nucleotide diversity (π) in comparison to randomly sampled genomic regions of the same size (Fig. 4A). Firstly, consistent with inversion polymorphisms that suppress recombination between haplotypes, the haploblock regions exhibited significantly lower recombination rates compared to the randomly sampled genomic regions. Secondly, these haploblock regions had a higher deleterious mutation load relative to the rest of the genome, consistent with heightened Hill-Robertson interference in inverted regions due to the suppression of recombination [37]. Lastly, the haploblock regions demonstrated significantly higher nucleotide diversity and genetic differentiation across populations compared to other genomic regions, further supporting the ‘local adaptation hypothesis’ [34–36]. Overall, while the haploblocks identified here show inversion-like characteristics, further independent evidence, such as long-read sequencing and comparative genetic mapping, is needed for additional confirmation.

**Fig. 4.**
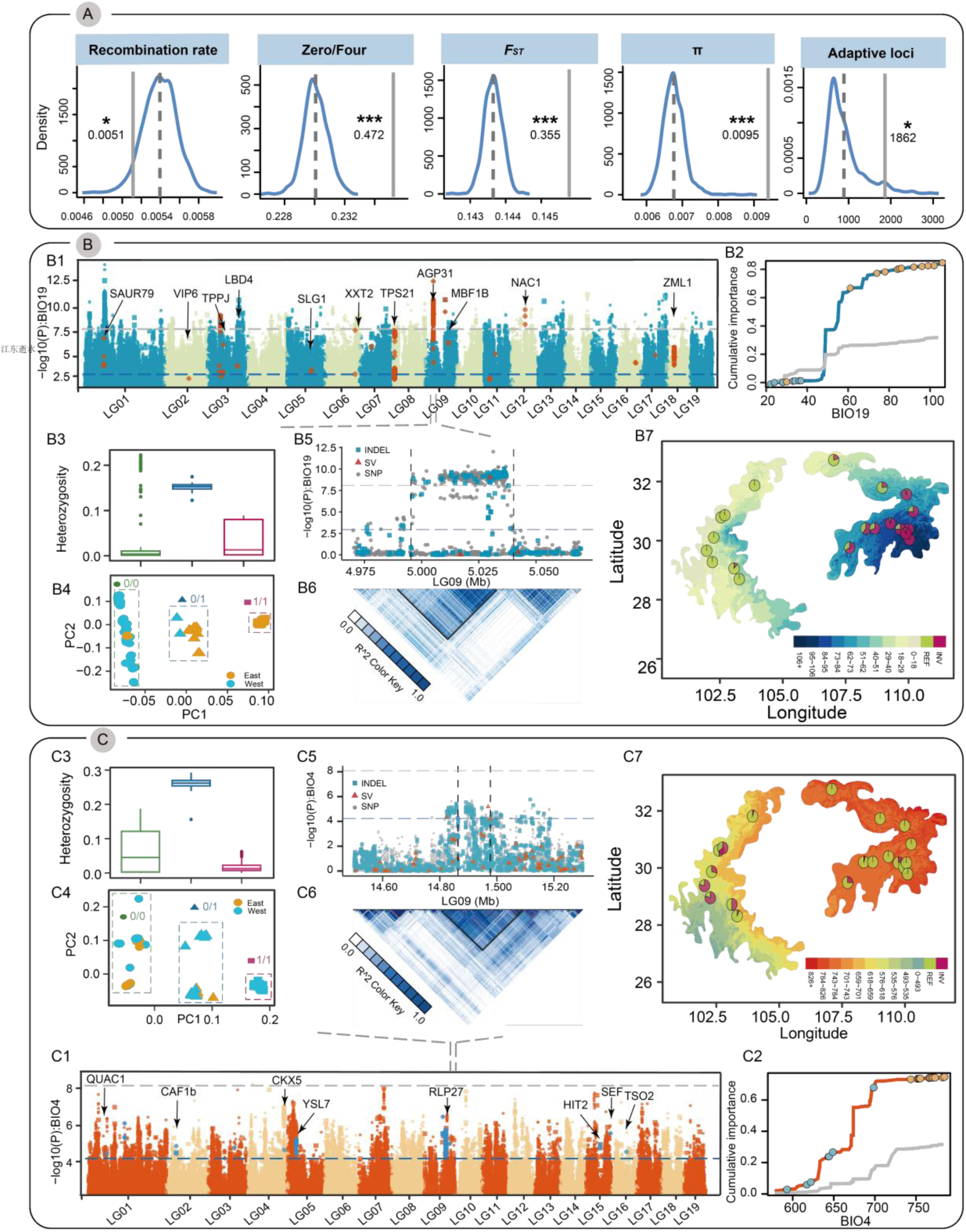
Haploblocks facilitate local adaptation. **(A)** Assessment of genomic features of candidate haploblocks. The diagram compares recombination rate, mutation load (ratio of 0-fold to 4-fold polymorphic sites), *F_ST_*, nucleotide diversity (π), and the number of adaptive loci between haploblocks (grey solid line) and permutation results from 1,000 randomizations (blue solid line). The dashed line represents the mean value across randomizations. Significance is assessed by permutation test and indicated by asterisks (*** *P*<0.001, ** *P*<0.01, * *P*<0.05, ns: *P*>0.05). **(B, C)** Two examples of haploblocks contributing to environmental adaptation: **(B1, C1)** Manhattan plots for variants associated with Precipitation of Coldest Quarter (BIO19) and Temperature Seasonality (BIO4). Red and blue circles represent adaptive variants within haploblocks. Selected candidate genes are labeled at their respective genomic positions. Dashed horizontal lines denote significance thresholds (blue: FDR correction 0.05; grey: Bonferroni correction, adjusted *P*=0.05). **(B2, C2)** Cumulative importance of allelic change along environmental gradients for neutral variants (grey line) and adaptive variants associated with BIO19 (blue line) and BIO4 (orange line). **(B3, C3)** Comparison of heterozygosity within the example haploblock among clusters assigned by PCA results. **(B4, C4)** Clustering of individuals using *k*-means based on PCA results for each haploblock. **(B5, C5)** Local magnification of the Manhattan plots around the selected haploblocks. **(B6, C6)** Linkage disequilibrium (LD) block for the example haploblock, shown as mean *r*^2^ for paired windows. **(B7, C7)** Allele frequencies of the example haploblock across 20 populations. Pie charts represent the genotype of the example haploblock based on clustering results from B4 and C4. Colors on the map represent environmental gradients across the distribution range.

To further investigate the role of these haploblocks in the local environmental adaptation of *P. lasiocarpa*, we compared the number of environmental adaptive variants within the haploblock regions to those in randomly sampled reference regions of the same size. Our analysis revealed a notable presence of 1,862 putatively adaptive variants in the haploblock regions, significantly exceeding expectations (Fig. 4A). This strongly suggest that these haploblocks, particularly the putative inversion polymorphisms, constitute a crucial source of genetic variation that facilitates local population differentiation and environmental adaptation [8, 38]. For instance, a haploblock on chromosome 9, spanning 44,913bp, contained 508 adaptive variants (447 SNPs, 60 Indels, and one SV), which were strongly associated with Precipitation of Coldest Quarter (BIO19) (Fig. 4B). This haploblock is characterized by high LD, and PCA revealed three distinct genotype clusters within the region, with higher heterozygosity in the central cluster. This pattern aligns with the expectation of inversion polymorphism, where the two extreme clusters represent homozygous genotypes for two distinct haplotypes, and the central cluster represents heterozygotes (Fig. 4B). Notably, four genes were identified in this region, including *Polas22852*, orthologous to *Arabidopsis AGP31*, known to be involved in vascular tissue function during defense responses, including responses to Methyl jasmonate and abscisic acid (ABA) treatments [39]. Another noteworthy example is a putative adaptive inversion spanning 21,893bp, which trapped 43 Temperature Seasonality (BIO4)-associated variants (38 SNPs, 4 Indels, and one SV) (Fig. 4C). Three genes were located within this region, including *Polas23357*, the ortholog of which (*RLP27*) was found to be involved in signal transduction and defense responses [40].

### The role of environmentally induced gene expression plasticity in rapid climatic adaptation

In addition to genetic changes susceptible to selection, we also explored the potential role of immediate plastic responses to environmental shifts. This approach is crucial for understanding whether phenotypic plasticity can lead to subsequent genetic adaptations. Since gene expression changes are highly responsive to environmental fluctuations and often serve as a link between the genome and the phenotype, we focused on gene expression levels as primary traits. Our aim is to clarify how the interaction between expression plasticity and evolutionary adaptation shapes rapid responses to environmental change [10–12]. To achieve this, we analyzed gene expression differences under both controlled and stress-inducing conditions (heat and waterlogging) for plants sampled from two populations with distinct temperature and precipitation-related environments (Figure 5A). These two populations were designed as West and East ecotypes for the downstream analysis.

**Fig. 5.**
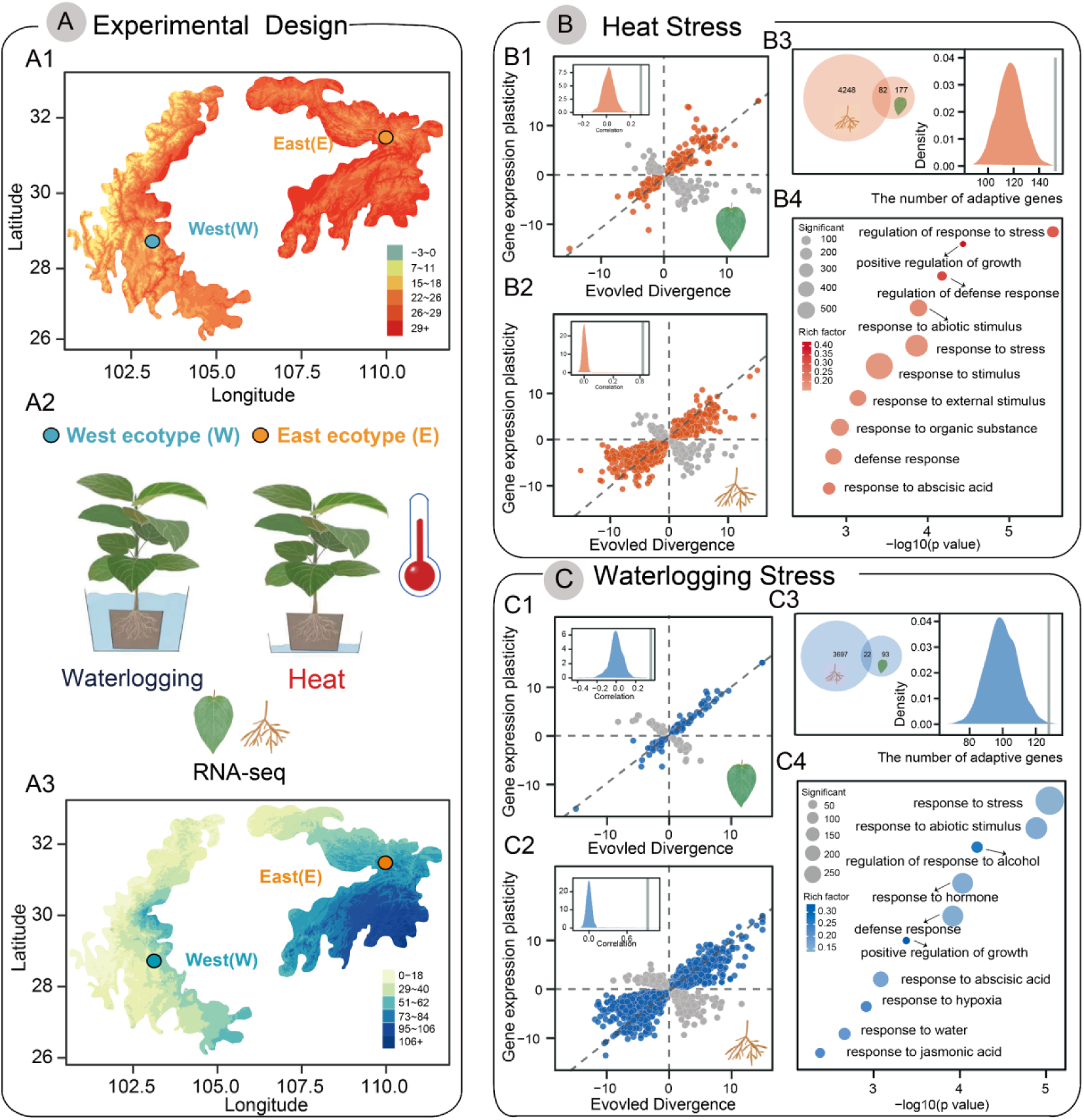
Plastic and evolved changes in gene expression. **(A)** Experimental Design: **(A1, A3)** Environmental gradients for Maximum Temperature of the Warmest Month (top) and Precipitation of the Coldest Quarter (bottom) across the distribution range of *P. lasiocarpa*. Circles on the map indicate the locations of the experimental individuals; **(A2)** Conceptual illustration of heat and waterlogging treatments, along with RNA-seq sampling of tissues. **(B, C)** Analysis of gene expression plasticity and evolved divergence under heat and waterlogging treatments: **(B1, C1; B2, C2)** Relationship between gene expression plasticity and evolved expression divergence for leaf (top) and root (bottom) tissues. Points represent differentially expressed genes (DEGs) showing both expression plasticity and evolved expression divergence. Orange and blue points indicate genes with a positive correlation between plasticity and evolved divergence (adaptive plasticity), while grey points denote a negative association (maladaptive plasticity). Insets show that the observed correlation coefficient (grey solid lines) is more positive than the permutation results; **(B3, C3)** Venn diagrams showing the overlap of adaptive plastic genes detected in leaf and root tissues under heat and waterlogging treatments, respectively. The figure on the right displays the number of genes with signatures of environmental adaptation, as evidenced by the GEA method, in the sets of adaptive plastic genes (grey solid line) compared to the permutation results; **(B4, C4)** Representative Gene Ontology (GO) enrichment categories of adaptive plastic genes in response to heat and waterlogging stresses.

We performed transcriptome sequencing on leaf and root tissues from both East and West ecotypes under two conditions: control and stress (heat and waterlogging), to simulate anticipated future environmental changes (Fig. 5A). Initial analysis using hierarchical clustering of expression profiles revealed that leaf and root tissue samples clustered into two distinct groups (Supplementary Fig. 14A). This finding was confirmed by principal component analysis (PCA), where PC1, accounting for 41.85% of the variance, reflected tissue-specific differences, and PC2, accounting for 12.92% of the variance, represented distinctions related to treatment types (Supplementary Fig. 14B). To further explore and compare the transcriptional responses of the two ecotypes to environmental stresses, we conducted pairwise contrasts between treatments within each ecotype (Supplementary Fig. 15A). We found that approximately 14% to 45% of the differentially expressed genes were shared between ecotypes, with variation depending on the environmental conditions (Supplementary Fig. 15B). Differentially expressed genes between treatments were enriched for GO terms associated with heat and water stress responses, respectively (Supplementary Fig. 15C).

To further investigate and characterize the relationship between plastic and evolved transcriptional responses to environmental changes, we compared the stress-induced gene expression plasticity in the West ecotype with the evolved expression differences between the West and East ecotypes (Supplementary Fig. 16A2, B2). The rationale behind this comparison assumes that the West ecotype represents the population akin to the “ancestral” state, whereas the East ecotype is perceived to be closer to the “evolved” state as it more closely aligns with and adapts to changing environments characterized by higher temperature and precipitation (Supplementary Fig. 16A1, B1). Therefore, if gene expression plasticity is adaptive, we would expect the direction of plastic responses to align with the direction of evolved responses. Conversely, if plasticity is maladaptive, we would anticipate that plastic responses would oppose evolved responses (Supplementary Fig. 16A3, B3).

We identified 1,078 genes whose expression significantly diverged between the West and East ecotypes under control conditions in leaf tissues (FDR<0.05). Among these, 259 genes exhibited heat-induced plastic responses in the same direction as evolved responses, while 154 genes showed oppositing expression directions (Fig. 5B1). Furthermore, there was a strong positive correlation between plasticity and evolved divergence (Spearman’s ρ = 0.33, Fig. 5B1, inset). To account for potential statistical artifacts from regression toward the mean, we conducted a randomization test to estimate a null distribution of correlation coefficients. This test confirmed that the observed correlation was significantly more positive than would be expected by chance alone (binomial exact tests *P* < 0.01, Fig. 5B1, inset). These results suggest that many of the expression responses to heat stress are adaptive, supporting the idea that adaptive plasticity contributes to local environmental adaptation. In root tissues, gene expression profiles showed a similar pattern, with significantly more adaptive plasticity under heat stress (Spearman’s ρ = 0.82, Fig. 5B2). Additionally, expression correlation coefficients between waterlogging-induced plasticity and evolution were 0.36 for leaf and 0.76 for root tissues, respectively (Fig. 5C1; Fig. 5C2), further reinforcing the pattern that plastic expression responses align with evolved responses.

Among the 4,507 genes displaying heat-induced adaptive plasticity (259 in leaf and 4,330 in root tissues), we observed a significant enrichment in genes associated with genetic adaptation to local environments, as previously identified (Fig. 5B3). This supports the role of selection on adaptive gene expression plasticity. GO enrichment analysis further revealed that these adaptive plasticity genes were significantly enriched in terms related to stress and defense responses, such as “regulation of response to stress”, “defense response”, and “response to abiotic stimulus” (Fig. 5B4). Notably examples include *Polas12092*, which is orthologous to *Arabidopsis ILL6* and involved in the jasmonic acid signaling pathway in response to abiotic stress [41]; *Polas02648*, orthologous to *SMXL7*, a negative regulator of drought resistance through regulation of the strigolactone signaling pathways [42]; and *Polas06886*, orthologous to *WRKY11*, which plays crucial roles in heat and drought tolerance in rice and *Arabidopsis* [43] (Supplementary Fig. 17A).

Similarly, among the 3,812 genes showing waterlogging-induced adaptive plastic responses (115 genes in leaf and 3,719 genes in root tissues), we also observed significant enrichment of genes associated with genetic adaptation (Fig. 5C3), reinforcing the role of selection on these gene sets. GO analysis indicated that genes with putatively adaptive plasticity were enriched in terms related to “regulation of response to alcohol,” “response to hypoxia,” “response to abscisic acid,” and “response to jasmonic acid,” highlighting their importance in waterlogging stress responses (Fig. 5C4). For example, several genes from key hormone- and stress-responsive transcription factor families (i.e. WRKY, ERF, and DOF), including orthologs of *CDF2*, *ERF1* and *WRKY40* (Supplementary Fig. 17B), are critical for plant survival and reproduction under various biotic and abiotic stresses [44, 45].

### Evolutionary genomic prediction of population-level vulnerability to future climate change

Accelerated climate change is profoundly posing a threat to global and local biodiversity, especially in mountainous areas that harbour exceptional endemism and species richness. The mountain regions around Sichuan Basin have experienced repeated and drastic climatic oscillation [46]. Consequently, our study offers a valuable opportunity to enhance our understanding of species’ response to future climate change in this crucial region. To explore the landscape of adaptive variation, we calculated an adaptive index, which estimates genetic similarities and differences based on the redundancy analysis (RDA), and mapped it across the geographic distribution of *P. lasiocarpa* [47]. After adjusting for correlations among environmental variables, we conducted RDA using the remaining seven environmental variables as predictors. This analysis revealed that most of the adaptive variation is concentrated along RDA1, which explains 78.83% of the variance. Extrapolating the adaptive gradients associated with RDA1 onto the landscape revealed a striking contrast in the adaptation index: eastern areas are characterized by warmer annual temperatures and higher precipitation during the warmest quarter (positive RDA1 scores), while western areas exhibit a strong shift in precipitation seasonality (negative RDA1 scores) (Fig. 6A).

**Fig. 6.**
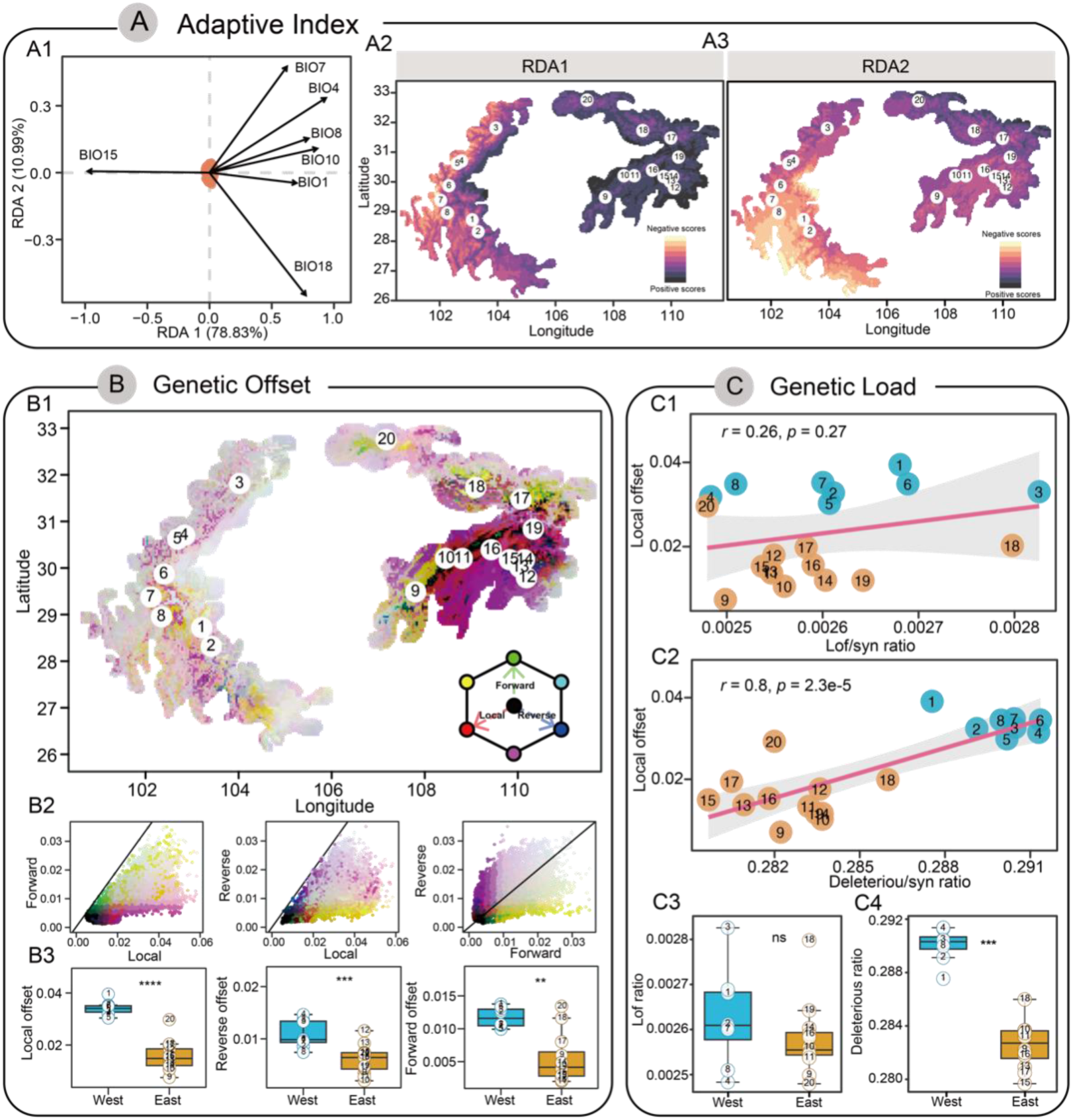
Genomic vulnerability prediction in response to future climate change. **(A)** Adaptive landscape across species range: **(A1)** RDA results showing the association between adaptive loci and the seven climatic variables. **(A2, A3)** Spatial projection of the adaptive index along RDA1 and RDA2 across the range of *P. lasiocarpa*. **(B)** Predicted genetic offsets to future climate change under SSP245 in 2061-2080: **(B1)** RGB map depicting local (red), forward (green) and reverse (blue) offsets. Circles represent the sampled populations. **(B2)** Bivariate scatter plots of the RGB map with 1:1 lines. Brighter cells (closer to white) indicate relatively high values for each offset, while darker cells (closer to black) indicate relatively lower values. **(B3)** Comparison of local, reverse, and forward offsets between western and eastern populations. **(C)** Relationship between genomic load and local genomic offset across *P. lasiocarpa* populations: **(C1, C2)** Relationship between two proxies of genetic load (y-axis: the ratio of derived deleterious/loss of function (Lof) variants to derived synonymous variants) and local genetic offset (x-axis). Blue and yellow circles represent western and eastern populations, respectively. The solid blue line denotes the best-fit linear regression line, with light-grey shading representing the 95% confidence interval. **(C3)** Comparison of genomic load between western and eastern populations. Significance was determined using the two-tailed Wilcoxon test, with significance indicated by asterisks above the boxplot (*** *P*<0.001, ** *P*<0.01, * *P*<0.05, ns: *P*>0.05).

Next, we investigate the capacity and vulnerability of natural populations to future climate change, aiming to predict which populations or regions might face the highest risks. To accomplish this, we utilized the Gradient Forest (GF) approach to quantify genomic offset, which assess the extent of genetic change required for a population to keep pace with future environmental conditions by modeling the turnover of adaptive genetic composition in response to projected future climates [48]. In addition to calculate local genetic offset, which assumes *in situ* tolerance, we further estimated two other metrics-forward and reverse genetic offsets-to assess the maladaptation of populations when simultaneously considering the contributions of *in situ* adaptation and migration to future shifting climates. Forward genetic offset calculates the minimum predicted offset, assuming populations could move (or be relocated) anywhere in Eurasia, while reverse genetic offset calculates the minimum offset between hypothetical populations in future climates and current populations within the existing range.

To measure the genetic changes needed for adaptation to new climate conditions across populations at various time points, we integrated projections from three different future climate models (CMCC-ESM2, GISS-E2-1-G, MPI-ESM1-2-HR) under moderate (SSP245) and severe (SSP585) shared socioeconomic pathways for the years 2080 (2061-2080) and 2100 (2081-2100), accounting for variability between climate models. The estimates of local, forward, and reverse offsets across different models showed highly consistent results (Supplementary Fig. 18), showing that populations in western areas exhibited greater genetic offset than those inhabiting eastern regions of the distribution ranges. Additionally, the spatial patterns of genomic offset align closely whether using all 19 climate variables or the seven uncorrelated variables identified from RDA analyses (Supplementary Fig. 19). Therefore, in the main text, we present the results averaged from the three climate models using all 19 climatic variables under the moderate scenario (SSP245) for the 2060-2080 period (Figure 6B). Again, the predicted patterns of substantially higher local, forward, and reverse offsets suggest that western populations distributed in the Hengduan Mountains are likely to be particularly vulnerable under future climates, as neither in situ adaptation nor migration are likely to be effective at reducing the population-level vulnerability to climate change in this region.

Finally, to further assess whether vulnerable western populations have already been adversely affected by recent unfavorable climate conditions, we measured genomic load. Deleterious genetic variants are expected to be less efficiently purged by purifying selection in populations already in decline coinciding with shifts in climate [49]. The genomic load for each population was estimated by calculating the proportion of derived loss-of-function (LoF) and deleterious variants relative to synonymous ones. Our findings reveal that the load of LoF variants did not show significant differences between western and eastern populations, implying that selection in both groups is effective at purging severely deleterious mutations. However, we observed a significant positive relationship between the ratio of deleterious to synonymous variants and the predicted local genomic offset across the sampled populations (Figure 6C). Western populations exhibited significantly higher loads of deleterious mutations compared to eastern populations, indicating that these populations may already be threatened by ongoing climate change. This shift has likely impacted the efficacy of removing moderately deleterious variants, contributing to detrimental effects in these regions.

## Discussion

Global climate change is increasingly exacerbating biodiversity loss. To better understand and predict species’ responses and dynamics in the face of climate change, it is essential to adopt an evolutionary perspective that incorporates both genetic and plastic effects [50]. Forecasts of future evolutionary responses to ongoing and future climate change rely heavily on the investigation of historical and contemporary adaptive evolutionary responses [3, 51]. In mountainous regions, highly heterogeneous environments and physical barriers often drive intraspecific genetic differentiation and local ecological adaptation in species [46]. Here we highlight the unique value of integrating evolutionary genomic approaches to predict and monitor species and population responses to global climate change, using *P. lasiocarpa*, an endemic and dominant tree species distributed in the biodiversity hotspot of the mountain regions of Southeast Asia. By leveraging a newly assembled high-quality reference genome and population genomic datasets, we explore the processes of adaptive evolution in response to climate change across historical, contemporary, and future timescales (Figure 7).

**Fig. 7.**
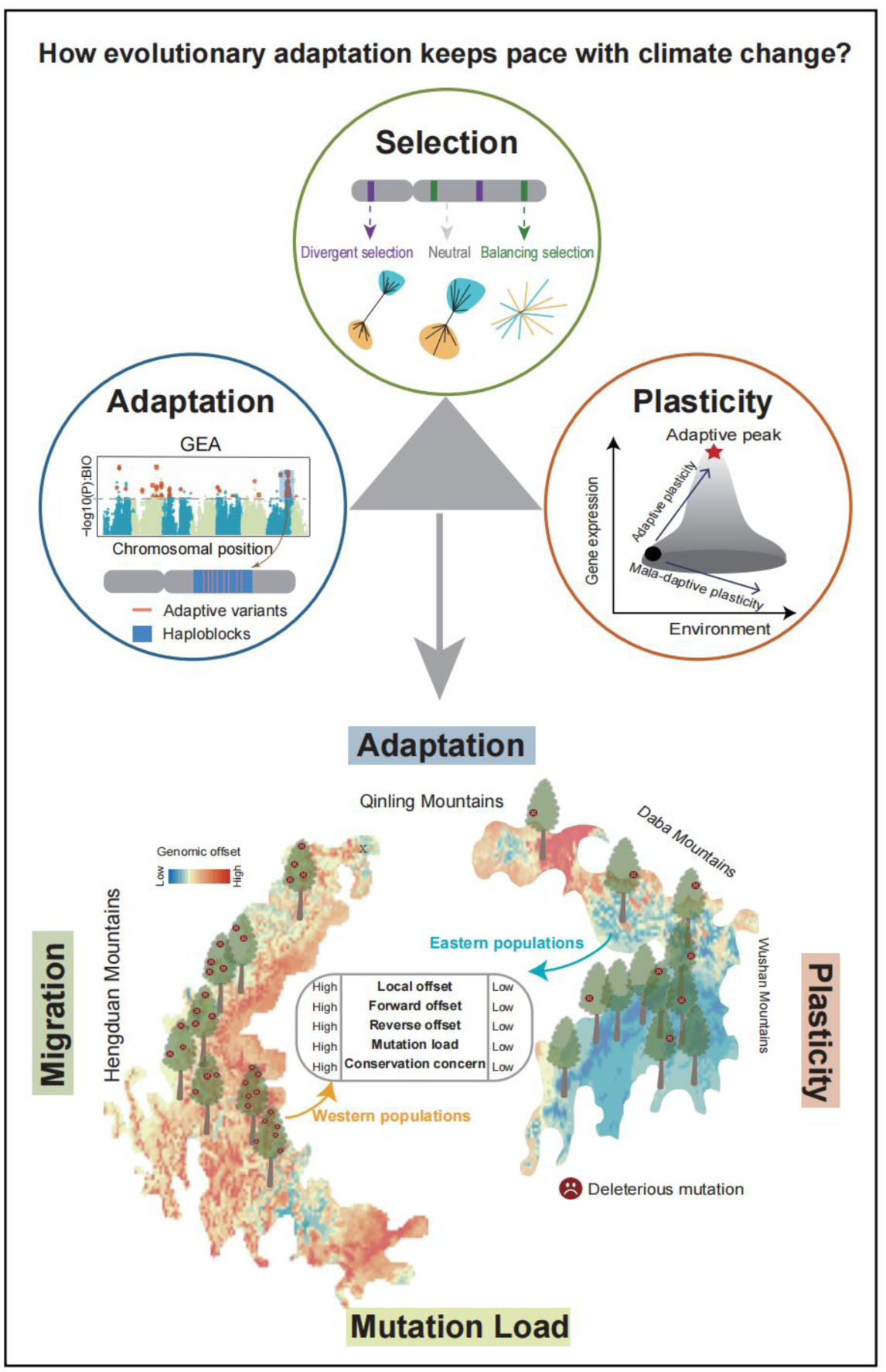
Schematic illustration of integrating evolutionary genomics to understand the rate of evolutionary adaptation in *P. lasiocarpa* populations in response to climate change. Large-scale population genomics, experimental evolution, and environmental modeling analyses were used in this study to elucidate the roles of evolutionary processes—such as selection, adaptation, and plasticity—in enabling populations to keep pace with climate change. These analyses highlight that western populations, primarily located in the Hengduan Mountains, are the most vulnerable to climate change and thus require priority in conservation and management efforts.

Our results first revealed a clear genetic separation between western and eastern populations, with the Sichuan Basin acting as both a geographical and genetic barrier, leading to a ring diversification pattern in *P. lasiocarpa*. Both lineages exhibited consistent demographic histories, suggesting that the climatic oscillations of the Quaternary similarly impacted both groups. Beyond neutral evolutionary processes like genetic drift and demographic changes, our findings demonstrated that genomic diversity and divergence were significantly shaped by divergent selection and long-term balancing selection. Regions under divergent selection, marked by strong signatures of positive selection in both lineages, exhibited higher levels of relative and absolute genetic divergence (*F*_ST_ and *d*_XY_) across populations and displayed a phylogenetic topology that clearly separated western and eastern lineages. Genes in these regions were enriched in categories related to environmental stimuli and stress responses, underscoring their crucial roles in driving differential adaptation to local climates in the two independent lineages [52]. Conversely, we identified regions with remarkable genetic diversity and elevated levels of trans-lineage shared polymorphism, likely maintained by long-term balancing selection [23]. These regions were significantly enriched for genes involved in metabolic processes and responses to biotic stresses. Together, these findings highlight the significant roles of divergent and long-term balancing selection, alongside historical demographic processes, in shaping the standing genetic variation to keep pace with climate change.

Identifying climate-associated adaptive genetic variation and characterizing its spatial distribution patterns can provide valuable insights into the adaptive potential of populations facing changing climates [2]. Our findings reveal numerous polymorphic haploblocks, characterized by features consistent with inversions, which significantly enrich locally climate-adaptive variants. Consistent with recent studies in sunflowers [8], deer mice [9], and an invasive weed species [53], our results suggest that haploblock polymorphisms may play a crucial role in maintaining the genetic basis of local adaptation. This is particularly important as the suppression of recombination within these regions can protect locally advantageous allele combinations from being diluted by migration, especially in the context of high gene flow [38]. However, we cannot yet determine whether the haploblocks detected here are associated with inversions or are closely linked alleles experiencing low recombination. Future research involving population-level long-read sequencing and molecular experiments is needed to better characterize these haploblocks. Additionally, we found that haploblock regions tend to accumulate deleterious mutations, likely due to the reduced efficacy of purifying selection caused by suppressed recombination [37]. This raises the unresolved question of the trade-off between the benefits of concentrating climate-adaptive variations and the costs of accumulating deleterious mutations within haploblocks. Addressing this paradox will require further research.

In addition to genetic responses to climate change, molecular phenotypic plasticity offers an immediate buffer against its negative effects. Our findings reveal that plastic gene expression changes generally align with evolved responses, indicating the predominance of adaptive plasticity to environmental stresses, as observed in multiple other systems [12, 54, 55]. Initial adaptive plasticity may serve as a critical stepping stone for populations to survive and persist under changing and novel environmental conditions. This short-term acclimation response can create more opportunities for natural selection to act, ultimately facilitating long-term adaptive evolution [56]. This process may be particularly important for rapid, within-generation responses to climate change in long-lived trees [57]. Indeed, we found that genes exhibiting adaptive expression plasticity show significant enrichment for climate-adaptive genetic variation, suggesting that genetic changes may have occurred in these genes. This might be because these populations have frequently encountered stressful environmental conditions, leading to gene expression plasticity being partly shaped to be adaptive under recurrent selective pressures, potentially resulting in genetic assimilation [58].

Finally, exploring and understanding the evolutionary forces and genomic architecture underlying contemporary climate adaptation provide a crucial baseline for forecasting species’ responses to ongoing and future climate change [2, 4]. By considering intraspecific adaptive variation and the mitigating effects of migration, our findings indicate that western populations are likely to be particularly vulnerable to future climate change, exhibiting higher local, forward, and reverse genomic offsets compared to eastern populations. Additionally, we found that western populations carry a greater genetic load of mildly deleterious mutations than eastern populations. While other factors affecting the demography of western populations cannot be ruled out, our results indicate that past and ongoing climate change may have intensified the decline of these populations, weakening purifying selection and reducing their fitness in changing environments. Furthermore, given the prominent role of adaptive gene expression plasticity in western populations, it remains uncertain whether this adaptive plasticity will weaken selection intensity, buffer genetic response, or constrain adaptation to altered environmental conditions [13]. Therefore, the interplay between plasticity, genetic variation, selection, and adaptation requires further investigation, particularly in understanding how plastic responses may either facilitate or impede evolutionary responses to climate change.

The western populations are primarily distributed in the Hengduan Mountains, a region known for its highly heterogeneous environments and significant biodiversity, with many endemic mountainous species [17]. Should these key forest tree species become vulnerable to future climate change, it could further jeopardize the local ecosystem and impact the entire endemic flora and fauna of the region. To mitigate local maladaptation, assisted gene flow from eastern to western populations could be an effective strategy, particularly given the natural genetic barrier created by the Sichuan Basin that separates these populations [59]. By introducing favorable genotypes from the eastern populations, which are preadapted to warmer and wetter climates, this approach may enhance the adaptive capacity of western populations to new climatic conditions. However, while our study integrates various evolutionary processes—including demographic histories, selection pressures, local adaptation, gene expression plasticity, and genetic load—to offer a comprehensive understanding of population-level vulnerability to climate change, further carefully designed common garden and transplant experiments are needed [60, 61]. These experiments will help validate our genomic forecasts and assess the practical benefits and risks of assisted gene flow for effective conservation and management strategies.

## Acknowledgements

This work was supported by the National Key Research and Development Program of China (2022YFD2201200). National Natural Science Foundation of China (32371695 and 31971567) and Fundamental Research Funds for the Central Universities (2023SCUNL105) to J.W..

## Author contributions

J.W. conceived and supervised the study. Z.L., Y.S., J.F., T.S. analyzed the data. J.F. and X.Z. conducted the stress experiments. K.M. and L.Z. handled the sampling and material collection. Z.L. and J.W. wrote the manuscript, with the input from L.R. and J.L.. All authors approved the final version of the manuscript.

## Data availability

All data needed to evaluate the conclusions in the paper are present in the paper and/or the Supplementary Materials. All sequencing data in this study will be deposited in National Genomics Data Center (NGDC) during reviewing process. All scripts used in this study will be available at https://github.com/jingwanglab/Evolutionary-genomics-of-Populus-lasiocarpa upon publication.

## METHODS

### Genome sequencing and assembly

Fresh young leaves of a wild *P. lasiocarpa* individual located in Xinqiao Town, Meishan City, Sichuan Province, China (103° 51’ 36”E, 30° 9’ 36” N) were harvested and frozen in liquid nitrogen. High-quality genomic DNA was extracted from leaves using the CTAB procedure [61]. For the long-read sequencing, libraries with >20 Kbp DNA fragments were constructed and sequenced on PromethION platform with Oxford Nanopore Technologies (ONT). For short-read sequencing, 150 bp paired-end libraries with an insert size of 350 bp were generated and sequenced on Illumina HiSeq X Ten platform. To achieve chromosome-scale genome assembly, a Hi-C sequencing library was constructed and prepared with Dpnll restriction enzyme, followed by sequencing on the Illumina NovaSeq platform.

We performed the *de novo* assembly process in the following steps. Initially, the long sequencing reads, after self-correction, were assembled into contigs using NextDenovo (https://github.com/Nextomics/NextDenovo). To obtain a high-quality assembly, polishing was performed with both Nanopore reads for three rounds using Racon[62] and Illumina reads for four rounds using Nextpolish v1.0.5[63]. Heterozygous contigs were then removed from the resultant contig-level assembly using purge_haplotigs v1.1.1[64] with parameter settings -l 5 -m 66 -h 170. Next, Hi-C reads filtered by fastp v0.20.0[65] were aligned onto the contig-level assembly using bowtie2 v2.3.2[66]. Each pair of the uniquely mapped reads was then merged to cluster, orientate, and link the contigs into 19 pseudo-chromosomes using LACHESIS[67]. Finally, to validate the completeness and accuracy of the assembly, we calculated the Benchmarking Universal Single-Copy Orthologs (BUSCO)[21] score and estimated the Illumina short reads mapping rate using BCFTOOLS v1.11[68].

### Gene prediction and functional annotation

Before gene prediction, preliminary transposable element (TE) annotation was performed by EDTA v1.9.3[69], and TEs annotated as “LTR/unknown” were re-classified using TEsorter v1.2.5[70]. The TE libraries were then used to mask the whole genome sequences using RepeatMasker v4.10[71]. To predict the protein-coding genes, we integrated three strategies including homology-based, RNA-seq-based, and *ab initio* prediction. First, we mapped the protein sequences of six species (*Populus trichocarpa*, *Populus euphratica*, *Salix brachista*, *Salix purpurea*, *Arabidopsis thaliana* and *Vitis vinifera*) to *P. lasiocarpa* genome by TBLASTN (ncbi-BLAST v2.2.28)[72], and parsed the resultant alignments for homology-based predictions using Genewise v2.4.1[73]. Then, assembled transcripts from RNA sequencing reads of leaf and stem tissues, obtained using both genome-free and genome-guided methods with Trinity v2.8.4[74], were aligned to the reference genome using PASA v2.4.1[75] to conduct RNA-seq-based prediction. Additionally, Augustus v3.3.2[76] was used for *ab initio* gene prediction, utilizing gene models predicted through both homology-based methods and RNA-seq data. Finally, EvidenceModeler v1.1.1[77] was employed to integrate all above gene models into a comprehensive gene set, which was subsequently refined through three rounds of PASA v2.4.1.

Gene functional annotation was conducted based on BLAST with 1e-5 E-value cutoff against 9 well-known public protein databases: the protein families database (Pfam), the NCBI non-redundant protein database (NR), the interproscan database, the KEGG database, the Swiss-Prot protein database, the Eukaryotic Orthologous Groups of proteins (KOG), the Clusters of Orthologous Groups of protein (COG), the Translated European Molecular Biology Laboratory (TrEMBL) database and the Gene Ontology (GO) database. The putative domains and GO terms of genes were identified using the InterProScan v5.32-71.0[78] program with default parameters.

### Genome resequencing, read mapping, and variant calling

A total of 200 *P. lasiocarpa* wild accessions were collected from 22 natural populations throughout the natural distribution of this species for whole-genome resequencing. Sampled individuals were at least 100m apart from each other within each population. Genomic DNA was extracted from fresh mature leaves using a DNeasy® Tissue Kit (QIAGEN). Illumina paired-end libraries of 150 bp in length were constructed and sequenced on Illumina NovaSeq 6000 platform with a target coverage of at least 20× per individual.

Prior to read mapping, raw sequencing reads were trimmed and filtered using Fastp v0.20.0. The parameter settings of ‘-f 5 –F 5 -t 5 –T 5’ were used to trim five bases from both the front and tail of paired-end reads. The parameters ‘-n 0 -q 20 –u 20’ were used to retain reads with no N base, a Phred quality greater than 20, and less than 20% unqualified bases. The quality of reads was assessed using FastQC v.0.11.7[79] before and after trimming. The remaining read pairs were then mapped to our high-quality reference genome with BWA-MEM[80] and sorted with SAMtools v.1.9[81] with default parameters. PCR duplicates were marked using MarkDuplicates tool from Picard v.2.18.21 (https://broadinstitute.github.io/picard/). For genomic variant identification, SNP and Indel calling was performed using Genome Analysis Toolkit (GATK v.3.8.1)[82] and its subcomponents HaplotypeCaller to obtain per-sample variants. Joint genotyping was then conducted using the GenotypeGVCFs tool with the option “-all-sites” to output both variant and non-variant sites. SNPs and Indels were filtered using VCFtools v0.1.16 [83], retaining bi-allelic variants with a genotype missing rate of less than 20%, after treating genotypes with read depth (DP) < 5 and genotype quality (GQ) < 10 as missing. SVs were called with DELLY v0.8.38[26], and those longer than 50 bp with precise breakpoints and a genotype missing rate of less than 20% were retained, after treating genotypes with DP < 5 and GQ < 10 as missing. Finally, SNPable (https://lh3lh3.users.sourceforge.net/snpable.shtml) was implemented to mask genomic regions in which reads were not uniquely mapped and the variants located in these regions were further filtered.

### Phylogenetic, Admixture, and Population Structure Analyses

To analyze population structure, we retained only SNPs with a minor allele frequency (MAF) greater than 5%. Highly correlated SNPs were excluded by performing LD-based SNP pruning using PLINK v1.90b6.18[84] with the parameter setting “--indep-pairwise 50 10 0.2” (any SNP with a correlation coefficient (*r*^2^) >0.2 with another SNP within the sliding windows of 50 SNPs and advancing steps of 10 SNPs across the genome was removed). A Neighbor-joining phylogenetic tree was constructed using PLINK and MEGA-X[85], and visualized in Figtree v1.4.4 (http://tree.bio.ed.ac.uk/software/figtree/). A principal component analysis (PCA) was also conducted to further evaluate the genetic structure using smartpca program in EIGENSOFT v.7.2.1[86]. ADMIXTURE v1.3.0[87] was employed to examine population structure across all individuals, with the number of population ancestries of predefined number of genetic clusters *K* set from 1 to 10. The most suitable number of ancestries was determined by the *K* value with the lowest cross-validation error. Additionally, to evaluate and validate the quality of SVs detected, we performed population structure analysis on the pruned SVs dataset using the same method and parameters.

### Genetic diversity, linkage disequilibrium (LD) and population differentiation

Intra-group nucleotide diversity (π) and inter-group genetic divergence (*F_ST_* and *d_XY_*) were calculated over 100 Kbp non-overlapping window after accounting for both the polymorphic and monomorphic sites using the program pixy v1.0.4.beta1 [88]. For LD decay evaluation, the squared correlation coefficient (*r^2^*) for all pairs of SNPs with MAF > 0.05 within a 200-Kbp window was computed and plotted using PopLDdecay v3.41 [89]. To further account for the influence of the number of individuals used in the calculation, we also randomly sampled 10 individuals from each group to calculate these parameters.

### Estimation of demographic history and divergence time

The ancient and more recent demographic history were estimated using pairwise sequentially Markovian coalescent model (PSMC v0.6.5-r67 [90]) and SMC++ v1.15.4 [91], separately. Assuming a mutation rate of 3.75×10^-8^ (mutations per site per generation) and complying with 15 years as the generation [92], the scaled results were then converted into years and effective population size (*Ne*). PSMC analyses were performed for each group of 10 selected individuals, with 10 bootstraps per individual, employing the parameter settings -N25 -r5 -p “4+25*2+4+6”. The median estimate across 10 individuals was used to represent the fluctuations in *Ne* for each group. To infer the more recent demographic history, VCF file with high-quality polymorphic SNPs was converted to SMC format using the vcf2smc command in SMC++ and then the *estimate* command was run on each group with default parameters. To estimate divergence times, vcf2smc and split commands in SMC ++ program were run on the VCF file for each group pair with default parameters.

To explore the pattern of gene flow in the phylogenetic context, the pruned common SNP set was used and *P. wilsonii* (belong to the section *Leucoides* same as *P. lasiocarpa*) was selected as an outgroup. Then, the migration events were inferred by TreeMix v. 1.13 [93] with the parameters ‘-bootstrap 5000 -m 0-5 (-m the migration event)’. The identity-by-descent (IBD) blocks among all individuals were detected using beagle v4.1[94] with the following parameters ‘window = 100,000; overlap = 10,000; ibdtrim = 100; ibdlod = 5; ibdcm=1e-5’.

### Identification of genomic regions under selection

To identify genomic regions with abnormal population structure that may be under selection and show heterogenous patterns of relatedness, we applied local PCA to 180 individuals from the western and eastern populations using the R package *lostruct* v0.0.0.9000 [25]. Depending on the LD decay and the need to capture meaningful selection signals, we selected a window size of 250 SNPs for this analysis. The *P. lasiocarpa* genome was first divided into 40,380 contiguous and non-overlapping windows, each containing 250 SNPs. Windows that exhibited unusually large sizes, exceeding 20 Kbp in physical length, were removed due to the likely influence of a high proportion of repetitive sequences. To measure the similarity of individual relatedness across windows, a Euclidian distance matrix was computed and visualized using a multidimensional scaling (MDS) transformation. Window with extreme MDS1 values (top 0.5% and bottom 0.5% of MDS1 values) were defined as outlier windows. We further performed PCA and constructed the Neighbor-joining phylogenetic tree for the top outlier regions, bottom outlier regions and background regions, respectively.

As the two sets of outlier windows exhibited phylogenetic clustering topologies consistent with signatures of divergent and balancing selection, we conducted the following analyses to further assess selection by comparing them with the remaining background portions of the genome: (1) intra-group nucleotide diversity (π) and inter-group genetic divergence (*F_ST_* and *d_XY_*) were calculated within and between groups of western and eastern populations using the program pixy v1.0.4.beta1; (2) Tajima’s D statistics were calculated using VCFtools in 10 Kbp non-overlapping windows separately for western and eastern populations; (3) a composite likelihood ratio statistic (CLR) based on the genome-wide site frequency spectrum (SFS) of polarized SNPs was computed using SweepFinder2 [95] with 2 Kbp non-overlapping windows separately for western and eastern populations; (4) a cross-population composite likelihood ratio (XP-CLR) score was calculated using XP-CLR package [96] with 1 Kbp non-overlapping windows to specifically test for the signals of divergent selection; (5) a beta statistic (β) was computed by BetaScan2 [97] with 1 Kbp non-overlapping windows based on the genome-wide SFS of polarized SNPs was used to specifically test for signatures of long-term balancing selection. Finally, the structures of candidate genes in candidate regions were exhibited by R package gggenes (https://github.com/wilkox/gggenes).

### Detection of environment-associated genetic variants

To screen for locally adaptive environmental variants, we extracted and downloaded a total of 29 environmental variables corresponding to the geographical coordinates of the population collection sites (Supplementary Table 14). These variables included nineteen climate variables related to temperature and precipitation, and water vapor pressure (kPa) in the period of 1970-2000, with 30sec resolution, all retrieved from WorldClim public database (https://www.worldclim.org). Additionally, four soil variables in 2017 at a depth of 5-15 meters with a 1000-meter resolution were downloaded from the SoilGrids website (https://soilgrids.org). Ultraviolet (UV) radiation variables in the period of 2004-2013 for each month, with a 15 arc-minutes resolution, were obtained from glUV dataset (https://www.ufz.de/gluv/) and subsequently transformed to five variables, including UV radiations for each of the four quarters and the entire year.

We retained common variants (MAF > 5%), resulting in a total of 5,315,353 SNPs, 687,429 indels and 27,980 SVs, and employed two approaches to identify environment-associated variants. First, we used latent factor mixed modeling (LFMM)[29], implemented in the R package LEA v3.2.0 [98], to search for significant associations between allele frequencies and each of the 29 environmental variables. This analysis was conducted with two latent factors, based on the optimal number of ancestry clusters inferred with ADMIXTURE acorss the 20 populations, to account for the influence of population structure. For each environmental variable, LFMM was run with 5 repetitions, including a burn-in of 5,000 cycles followed by 10,000 iterations. The median of the resulting z-scores was calculated across the five runs, recalibrated by lambda, and the resulting *P-values* were adjusted for multiple testing using the false discovery rate (FDR) method, with an FDR of 5% set as the significance threshold. Furthermore, we employed a linear mixed model approach using the program GEMMA v 0.98.3 [30] to perform genome-wide environmental association studies, treating the environment as phenotype data. This method utilized a centered kinship matrix (-gk 1) generated from all 6,030,762 variants to account for relatedness among individuals and population structure. Similar to LFMM, we applied an FDR < 5% to control for multiple testing across the variant dataset.

Finally, to assess the reliability of the identified adaptive variants, we evaluated allele frequencies turnover along the environmental gradient and compared the patterns of these changes between the identified adaptive variants and a set of neutral references with the same number of variants. Following the method used in Aguirre-Liguori et al. 2021 [4], this analysis was conducted using machine learning algorithms implemented in the R package ‘gradientForest’ [99] to detect non-linear change in allele frequencies (response variables) along the climate gradient (predictor). We ran gradient forest with parameters “ntree=400, maxLevel= log2(0.368*n/2), corr.threshold=0.40” and calculated the cumulative importance of allele frequency changes as the sum of the split importance along climatic variables. The allele frequencies turnover along the environmental gradient was visualized by R package ggplot2. Additionally, to further evaluate the impact of geography and environment on shaping genomic variation of adaptive and neutral variants, we used linearized *F*_ST_ (*F*_ST_ /1-*F*_ST_) to investigate isolation by distance (IBD) and isolation by environment (IBE) through Mantel and partial Mantel tests, respectively, with Euclidean distance based on all scaled environmental variables used as the measure of environmental distance.

### Characterization of haploblocks

To identify genomic regions displaying abnormal population structure potentially generated by haploblocks or inversions, we conducted local PCA analysis using R package “lostruct”. This analysis was conducted with 60 SNP-wide windows (averaging 4,642bp) for each chromosome, aiming to capture smaller, more precise regions that could be grouped as haploblocks in subsequent analyses. We calculated Euclidean distances between PCA maps (using the first two principal components) to assess relatedness among windows and visualized these distances using MDS scores plotted against the positions of each window on the chromosome. Three-dimensional MDS axes were utilized to improve the detection of potential inversions. We defined genomic regions with at least ten adjacent windows having a z-score >1.5 as preliminary candidate haploblocks.

To provide further evidence for MDS outliers potentially caused by haploblocks, we conducted additional analyses to verify the following genomic signatures: (1) PCA of all SNPs within the candidate region should reveal three distinct clusters corresponding to 0/0, 0/1, and 1/1 genotypes; (2) the 0/1 genotype should exhibit the highest heterozygosity compared to 0/0 and 1/1 genotypes; and (3) high linkage disequilibrium (LD) should be observed within the haploblock region due to recombination suppression.

We then compared π, inter-population *F_ST_*, the ratio of zero-fold to four-fold synonymous variants, the number of climate adaptive loci and population-scaled recombination rate (calculated by the Interval program of LDhat [100] with parameter settings“-its 1000000 -bpen 5 -samp 2000”) between haploblock regions and randomly sampled genomic regions of the same size. Finally, we performed GO enrichment analysis using topGO v2.42.0 [101] to determine whether genes in haploblock regions were enriched for specific biological functions.

### Stress experimental design and RNA sequencing

Mature seeds were collected from the individuals of the West (103° 7’ 12’’ E, 28° 43’ 12’’ N) and East (109° 28’ 48’’ E, 29° 59’ 24’’ N) ecotypes of *P. lasiocarpa*. The seeds were first germinated in petri dishes and then transferred to pots, where they were grown in a greenhouse for four months under a temperature regime of 25 °C (day) and 20 °C (night), with a 16-hour photoperiod and an 8-hour dark period. After this growth period, the seedlings from both ecotypes were divided into three groups: control, heat-stress and waterlogging-stress, with each group consisting of three biological replicates. The control group was maintained under standard conditions of 25 °C/20 °C (day/night). For the waterlogging stress group, plants were transferred to tanks and subjected to full waterlogging, with the water level 3-5cm above the soil surface, for 15 days at 25°C/20°C (day/night). The heat stress group was placed in an incubator set to 42 °C (day) and 35 °C (night) for three days. On the final day of treatment, leaf and root tissues were collected from consistent positions across all replicates, flash-frozen in liquid nitrogen, and stored at -80°C. Total RNA was then extracted from these tissues, and RNA-seq libraries were prepared and sequenced on the DNBSEQ-T7 platform, generating 150 bp paired-end reads.

### Gene expression analyses

After filtering and trimming RNA reads using Trimmomatic v0.39 with default parameters, we mapped the clean reads to the reference genome using HISAT v2.2.1[102], and gene expression was quantified with StringTie[103]. The gene expression values were then normalized by Python script prepDE.py to generate reads count data. Based on this normalized data, we first created a correlation matrix and conducted PCA among the treatment samples and replicates. Genes with fewer than 10 reads were removed, and only the expressed gene were retained for downstream analyses.

To quantify expression changes, DESeq2 [104] was used to test for differential expression between control and stress-treated individuals in each group of the two ecotypes in both leaf and root tissues. Differentially expressed genes (DEGs) were identified using cutoff threshold of FDR < 0.05 and -2<log_2_ Fold Change>2. To assess whether plastic changes in gene expression under stress conditions correlated with evolved expression divergence, we compared plastic expression changes in the West ecotype (considered more “ancestral” given its greater distance from future climate conditions) to evolved expression changes between the West and East ecotypes (the logic underlying this comparison is detailed in Results section).

We first identified genes differentially expressed between the West and East ecotypes under control conditions. For these genes, we further compared the gene expression fold changes between control and stress-treated conditions in the West ecotype to the fold change in gene expression between the West and East ecotypes (Supplementary Fig. 16). To assess the correlation between plasticity and evolved divergence, we calculated Spearman’s rank correlation coefficient (ρ) using the values of log_2_ Fold Change. Given that the comparison involved two ratios sharing a common denominator, we further accounted for potential statistical artifacts by employing a randomization and resampling procedure to generate a null distribution of correlation coefficients. The candidate gene set was randomized 10,000 times, and the correlations between plastic and evolved divergence in these permuted results were used to create a distribution for comparison to the observed correlation. Finally, genes were categorized as exhibiting adaptive plasticity if the direction of expression plasticity aligned with evolved divergence, and as maladaptive if it was opposite. We then performed Gene Ontology (GO) enrichment analysis for the adaptive plastic genes.

### Adaptive index

To investigate the adaptive genetic landscape of *P. lasiocarpa,* we performed a redundancy analysis (RDA) using the R package “vegan v2.6-4” [105] with the setting of *K*=2 to calculate the adaptive index. Only LD-pruned adaptive variants and seven uncorrelated (pairwise correlation coefficients |r| < 0.7) environment variables (i.e. BIO1, BIO4, BIO7, BIO8, BIO10, BIO15 and BIO18) were utilized. The adaptive index was then projected across the species’ range to visualize geographic patterns of adaptation by R package ggplot2.

### Genomic offset assessment

For each sampling location, future BIOCLIM data for 19 climate variables at a 2.5 arcmin resolution for the periods 2061-2080 and 2081-2100, under two shared socioeconomic pathways (SSP245 and SSP585), were downloaded from the WorldClim CMIP6 website. Data from three climate models (CMCC-ESM2, GISS-E2-1-G and MPI-ESM1-2-HR) were integrated. A machine learning approach, Gradient Forest, was used to predict the genomic offset in response to environmental shifts. The GF model was built with parameters ntrees=500, maxLevel=log2(0.368*n/2), and corr.threshold=0.5, based on 19,955 adaptive variants and 15 environment variables that had identified adaptive variants.

Three metrics of potential maladaptation were calculated: local offset (population vulnerability *in situ* to new climate conditions), reverse offset (potential maladaptation if individuals could be relocated anywhere within the species range), and forward offset (minimum maladaptation assuming individuals could be moved anywhere within Eurasian continent) [47]. These metrics were visualized as the red, blue and green blue bands of an RGB image, respectively. Additionally, we assessed the variation in forward offset across different maximum allowable migration distances, including 100 km, 250 km, 500 km, 1000 km, and unlimited across the Eurasian continent. The results suggested that forward offset decreased substantially with increasing migration distances, stabilizing at 500 km and beyond (Supplementary Fig. 20). Therefore, forward offset results with unlimited dispersal distance were presented and projected on the map.

### Assessment of genetic load

To assess the deleterious genetic load in the *P. lasiocarpa* population, variants were annotated as either synonymous or non-synonymous, and SIFT score were calculated for the non-synonymous variants using SIFT 4G annotator [106]. Variants were then classified into four categories: loss of function (LOF), deleterious (SIFT score <0.05), tolerated (SIFT score >0.05), or synonymous. Additionally, the derived versus ancestral allelic state at each SNP position was determined using est-sfs [107], with *P. trichocarpa* as the outgroup. The ratio of the number of derived mutations (LOF and deleterious) to the number of synonymous variants served as used as a proxy for genetic load.

