## Supplementary_Files for "Evolutionary genomics predicts adaptive genetic and plastic gene expression responses to climate change in a key alpine forest tree species"

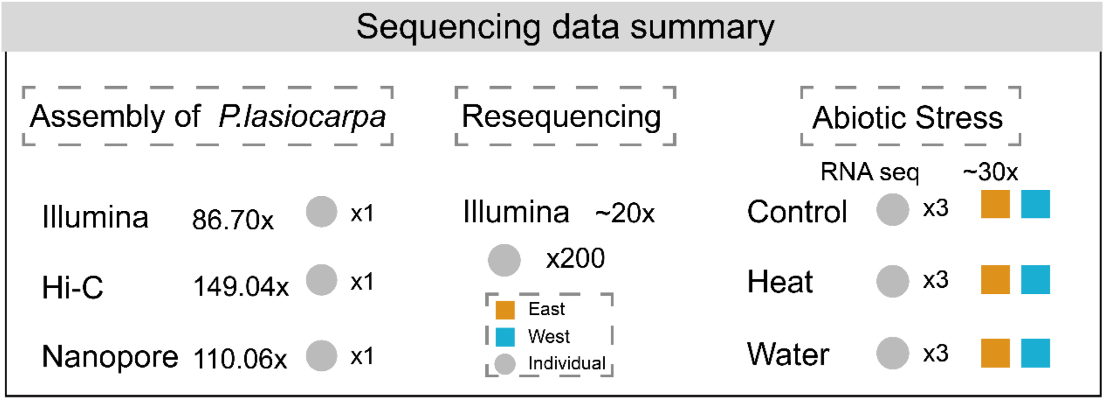


**Supplemental Figure 1.** Summary of sequencing data utilized in this study: including a high-quality genome assembly of *P. lasiocarpa*, re-sequencing data of 200 individuals across 22 populations, and RNA-seq data collected under control and stress conditions from two ecotypes.

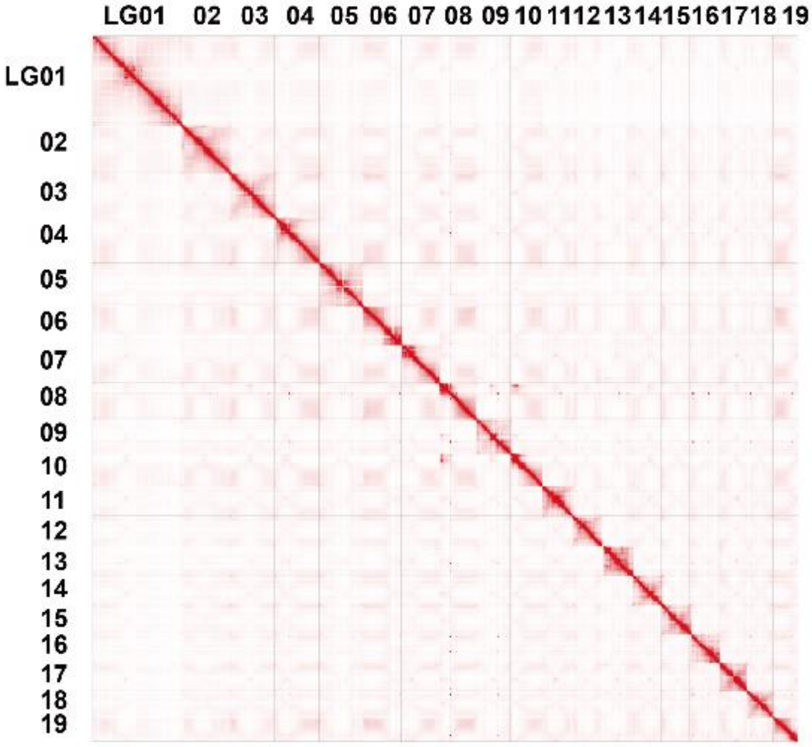


**Supplemental Figure 2.** Hi-C chromatin interaction map for the 19 pseudo-chromosomes of *P. lasiocarpa* genome.

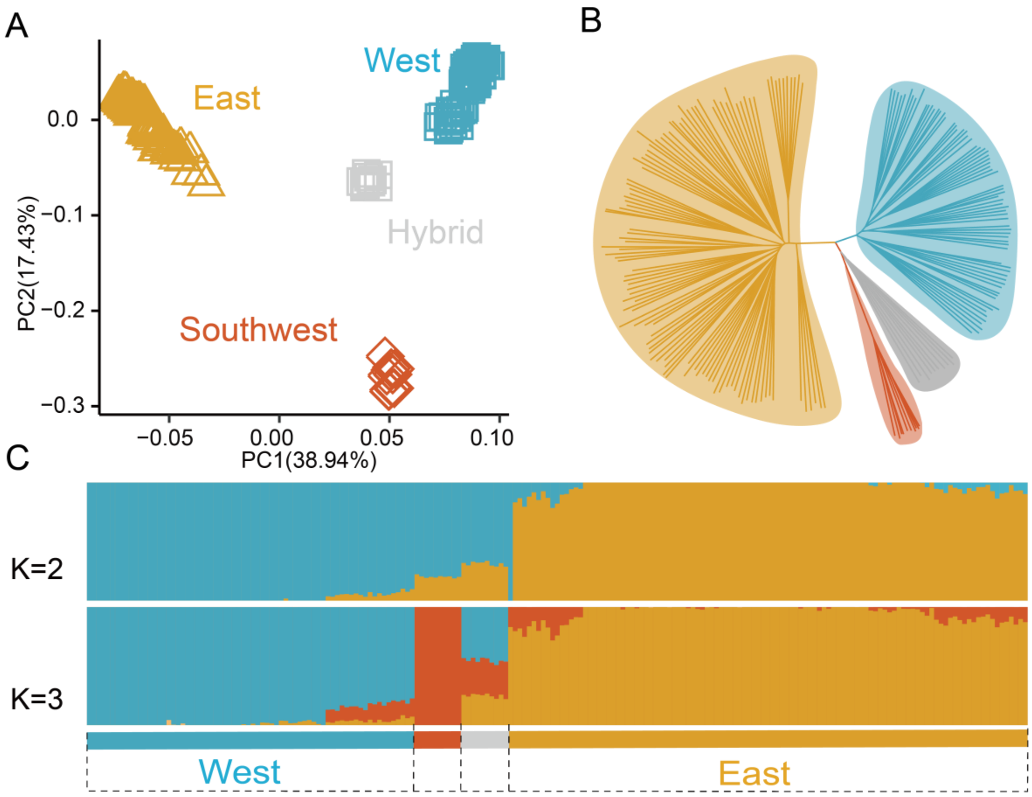


**Supplemental Figure 3.** Phylogeny and population structure of *P. lasiocarpa* based on the dataset of structural variations (SVs).

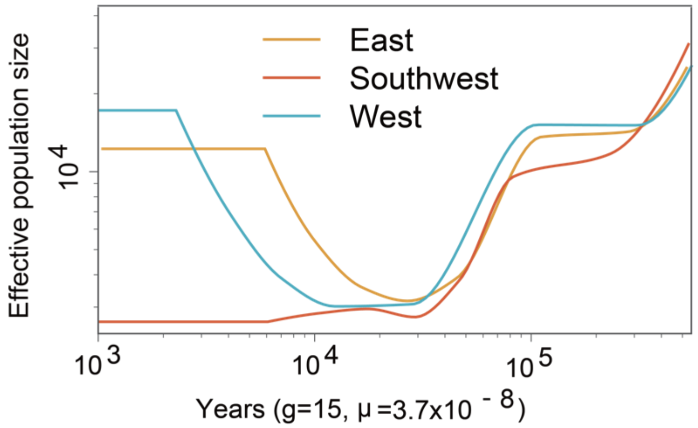


**Supplemental Figure 4.** Inferred the more recent effective population size over time using SMC++.

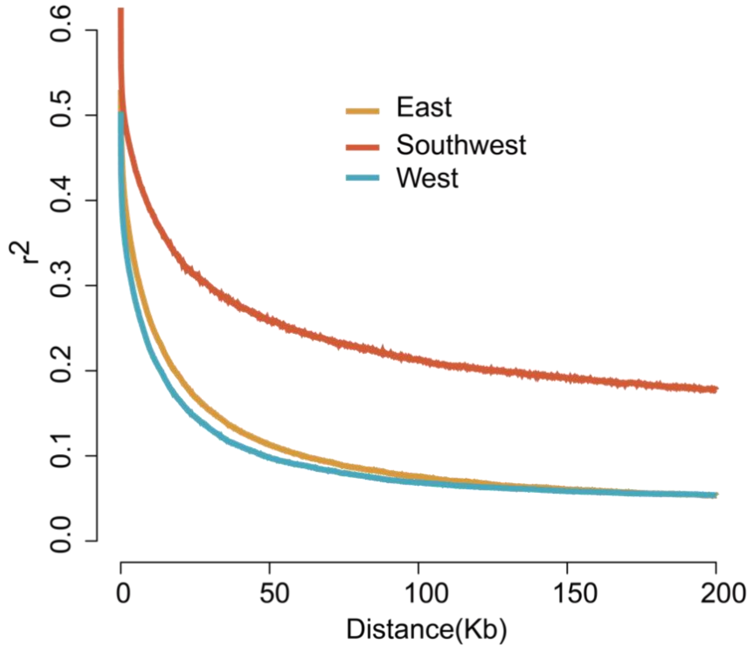


**Supplemental Figure 5.** Linkage disequilibrium decay in East, Southwest and West clusters.

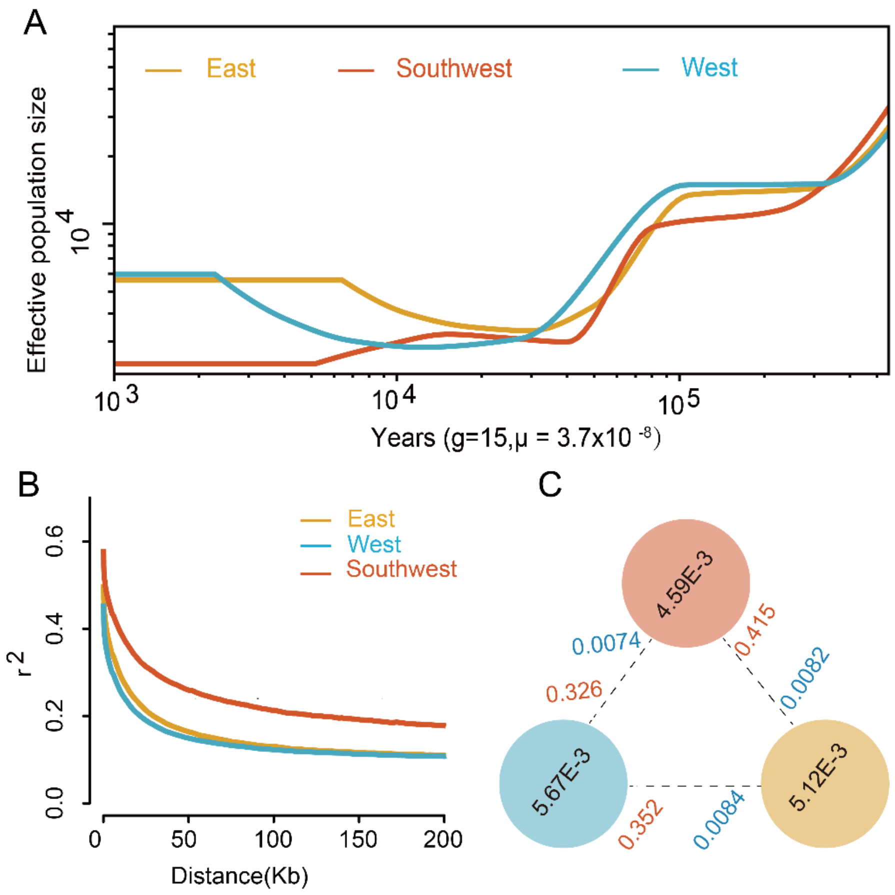


**Supplemental Figure 6.** Patterns of genomic diversity of *P. lasiocarpa* using the same number of individuals (randomly selecting 10 individuals) for the three different groups. **(A)** More recent demographic history revealed by SMC++**. (B)** Linkage disequilibrium decay estimated for the three groups. **(C)** Nucleotide diversity (π), Wright's fixation index ($F_{ST}$) and absolute genetic divergence ($d_{XY}$) (red: Southwest; blue: West; orange: East).

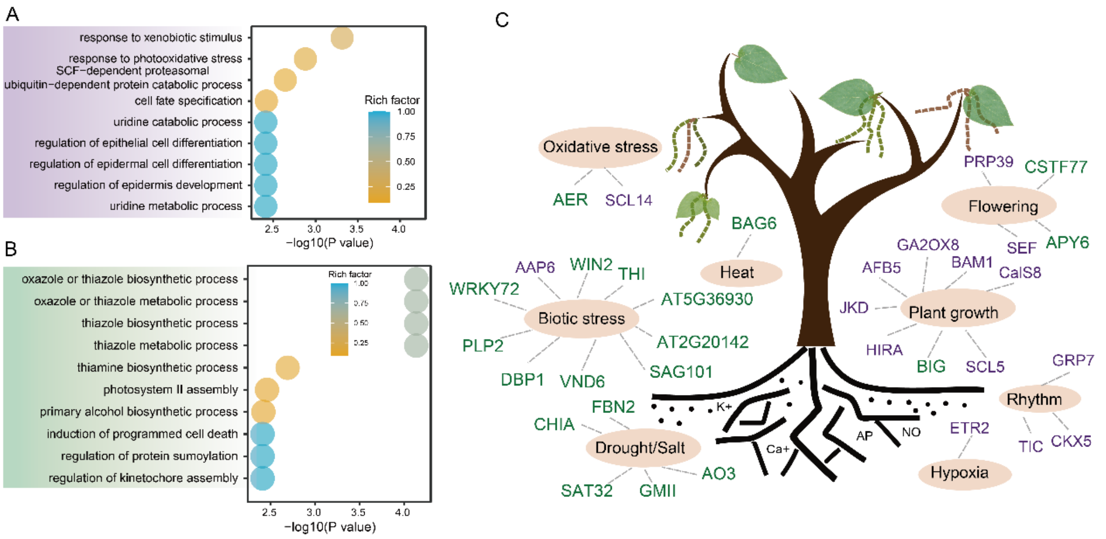


**Supplemental Figure 7.** Gene Ontology (GO) functional enrichment analysis of the candidate genes within the divergent selection regions **(A)** and balancing selection regions **(B)**. **(C)** The orthologous gene to *Arabidopsis* of candidate genes in divergent (colored by purple) and balancing selection (colored by green) regions.

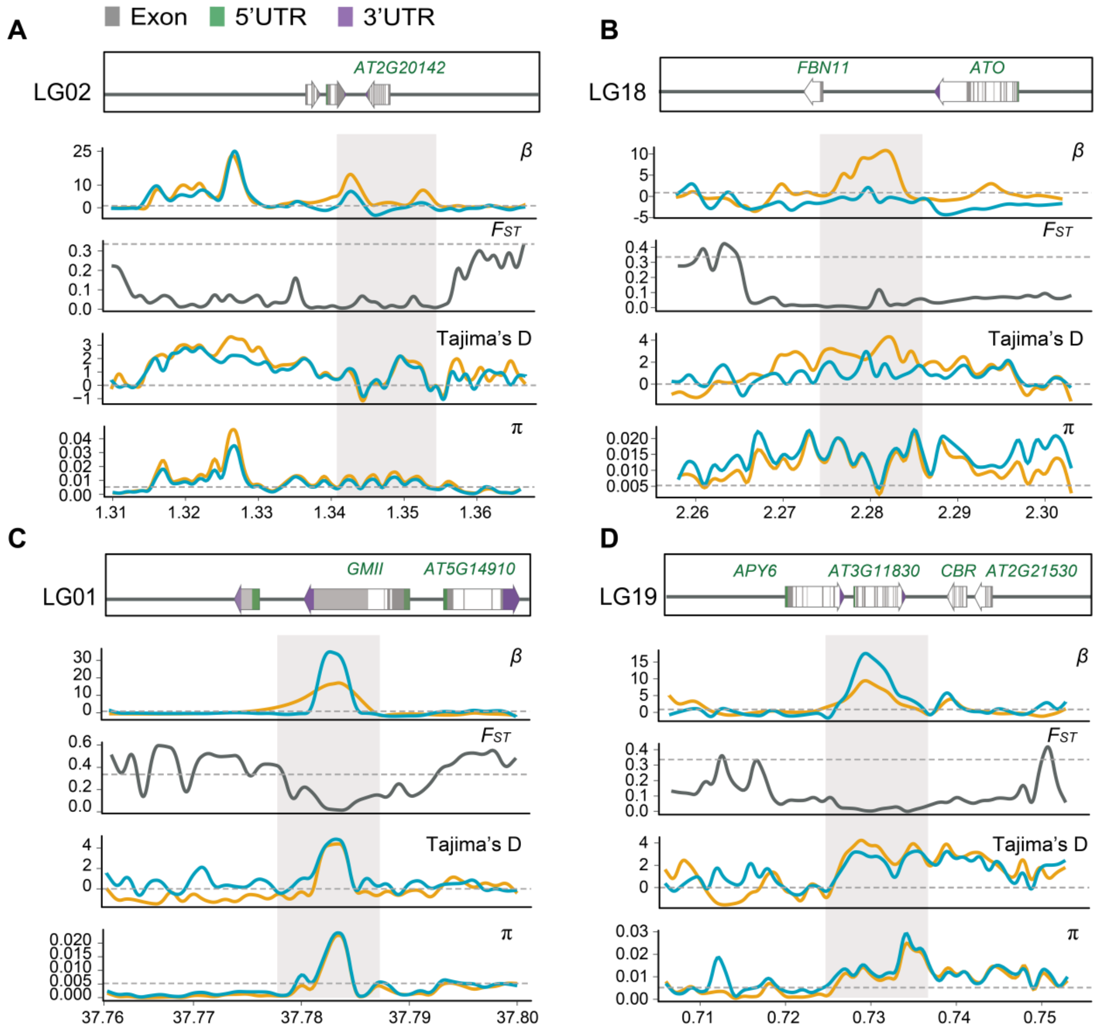


**Supplemental Figure 8.** The distribution of the statistics of Beta ($\beta$), $F_{ST}$, Tajima's D and $\pi$ around the example candidate genes within the balancing selection regions (gray shadows). Grey dashed lines represent the genome-wide significance level of the top 0.5% for the corresponding parameters. The structure of candidate gene is showed in the upper box.

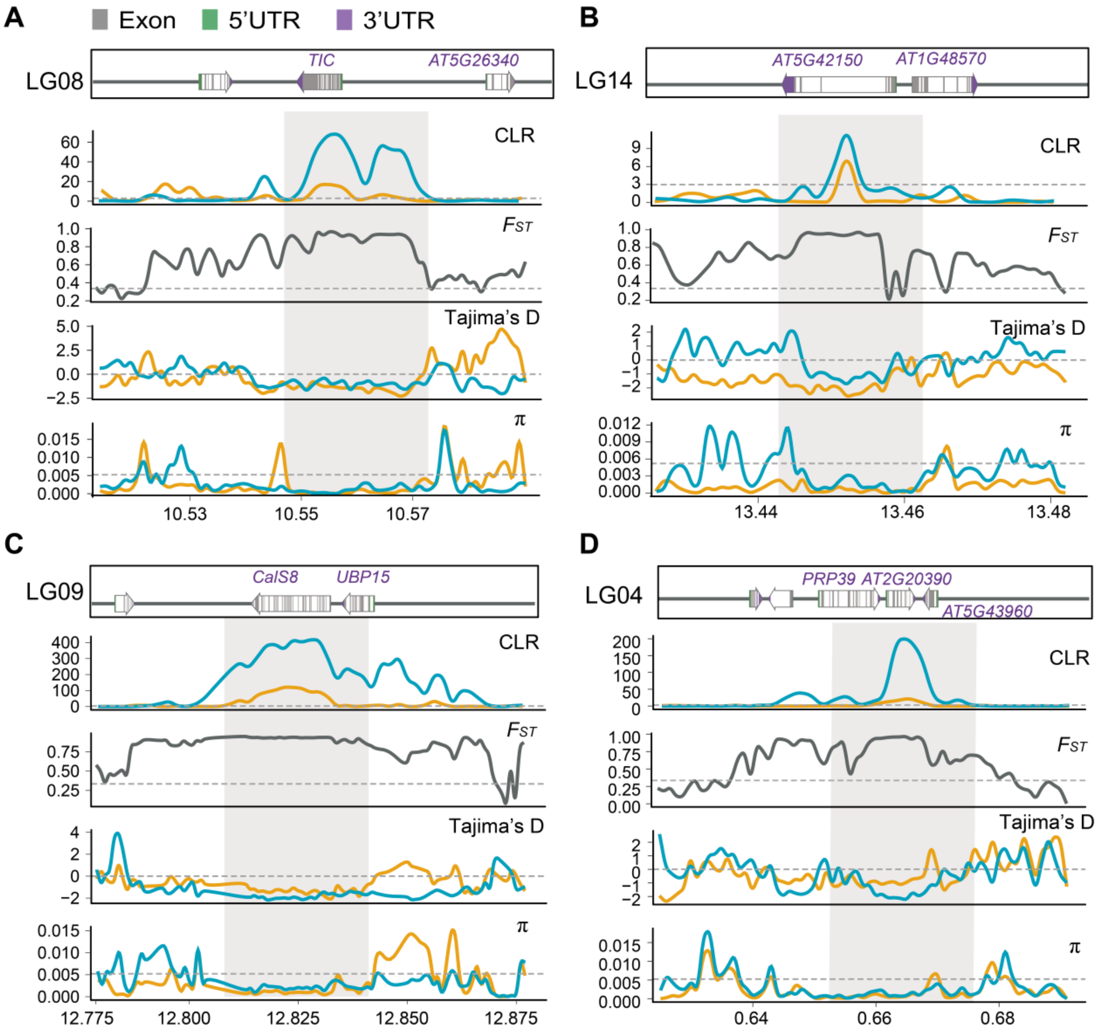


**Supplemental Figure 9.** The distribution of XP-CLR, $F_{ST}$, Tajima's D and π around the example candidate genes within the divergent selection regions (gray shadows). Grey dashed lines represent the genome-wide significance level of the top 0.5% for the corresponding parameters. The structure of candidate gene is showed in the upper box.

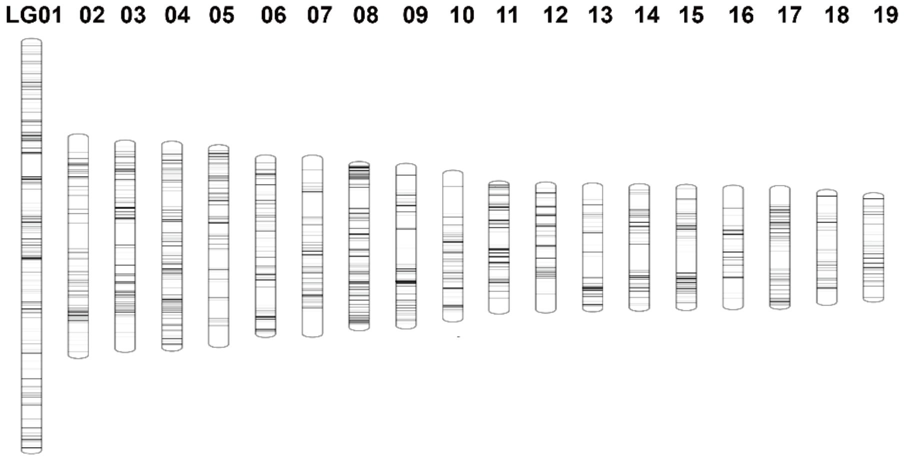


**Supplemental Figure 10.** Mapping of the location of the 19,995 environmentally adaptive variants identified by both LFMM and GEMMA across the 19 chromosomes.

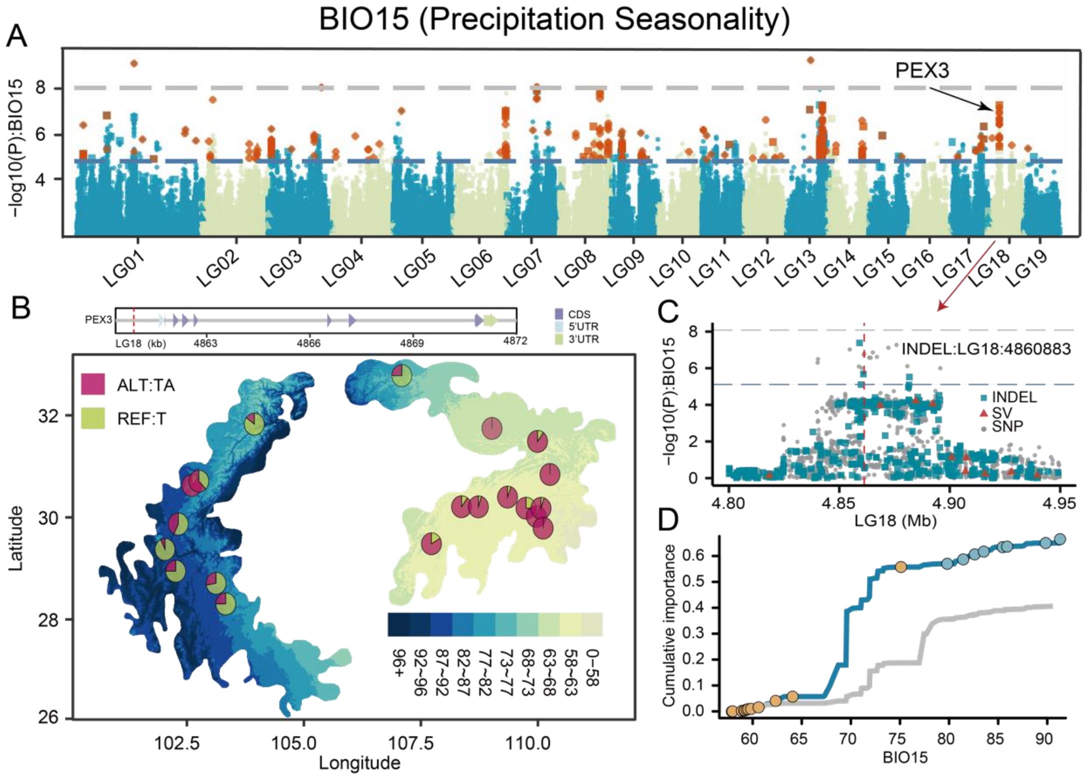


**Supplemental Figure 11.** Examples of adaptive variants of SNPs and short Indels. **(A)** Manhattan plot showing variants associated with specific environmental variables. Dashed horizontal lines represent significance thresholds (blue represents the FDR correction 0.05; grey line represents the Bonferroni correction, adjusted *P value* =0.05). The selected candidate gene is labeled at the corresponding genomic position. **(B)** Allele frequencies of a variant associated with an environmental variable across 20 populations. The map colors represent climate gradients across the distribution range under the current scenario. The structure of the candidate gene associated with this variant is shown in the upper box. **(C)** Local magnification of the Manhattan plots around the selected genes. **(D)** Cumulative importance of allelic changes along environmental gradients for neutral variants (grey line) and adaptive variants associated with specific environmental variables (colored line).

**Continuation Supplemental Figure 11.**

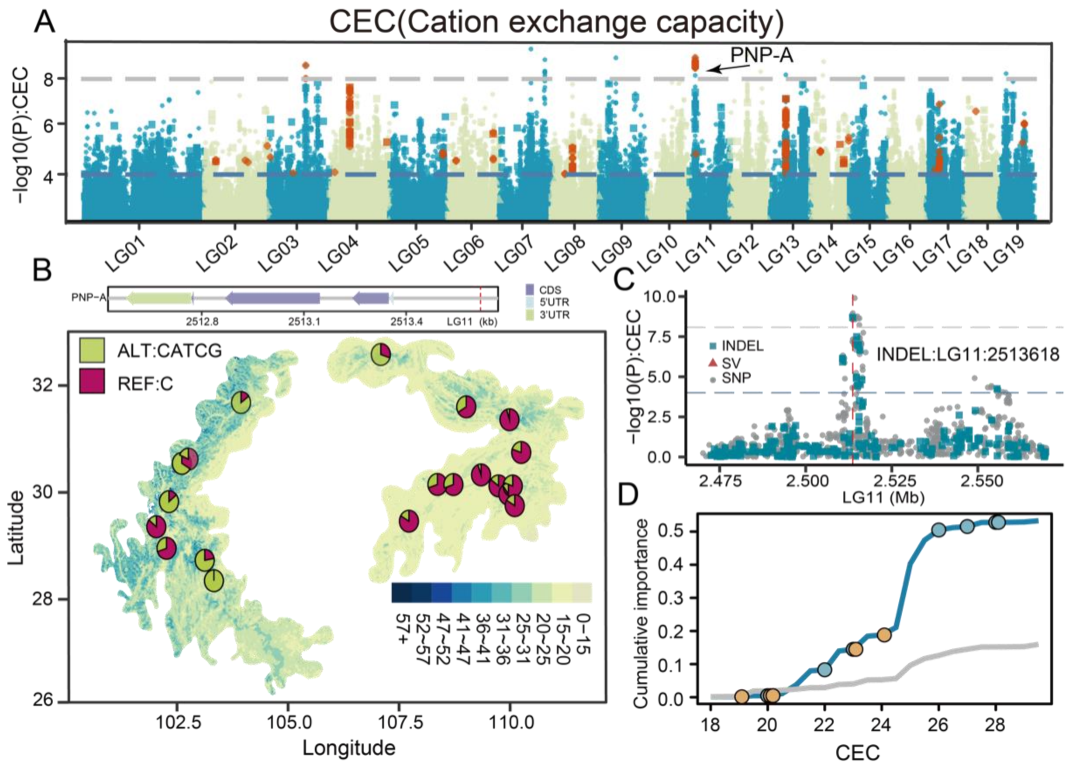

**Continuation Supplemental Figure 11.**

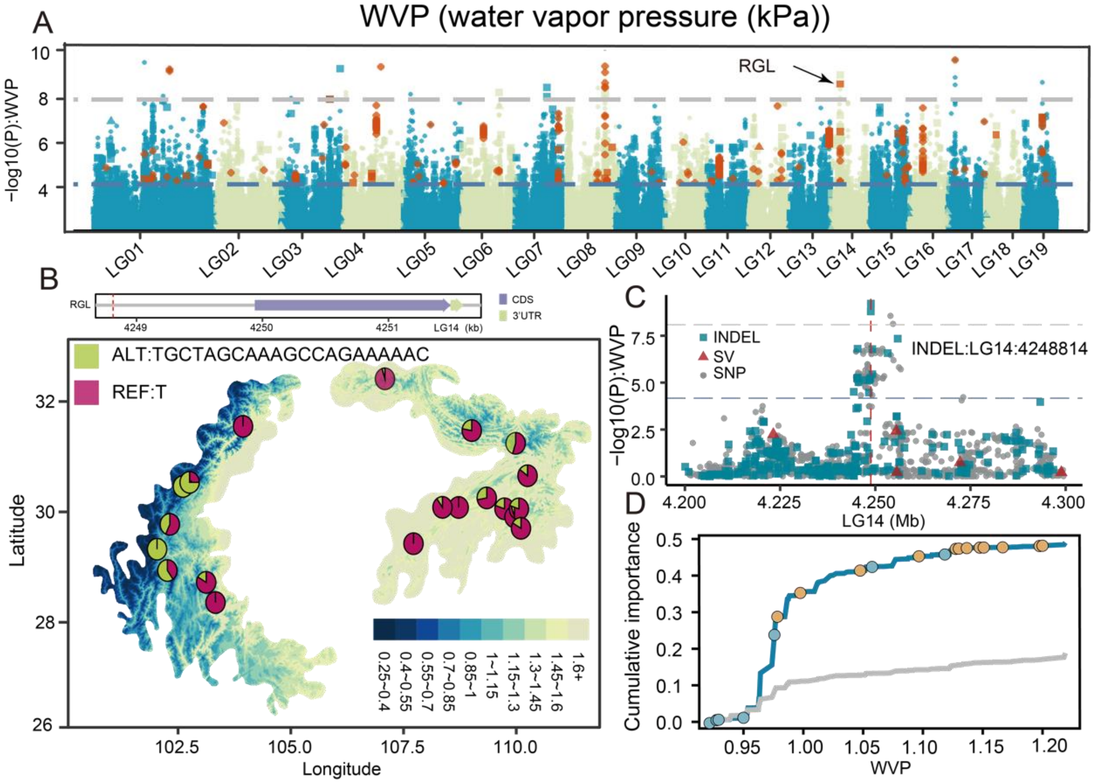

**Continuation Supplemental Figure 11.**

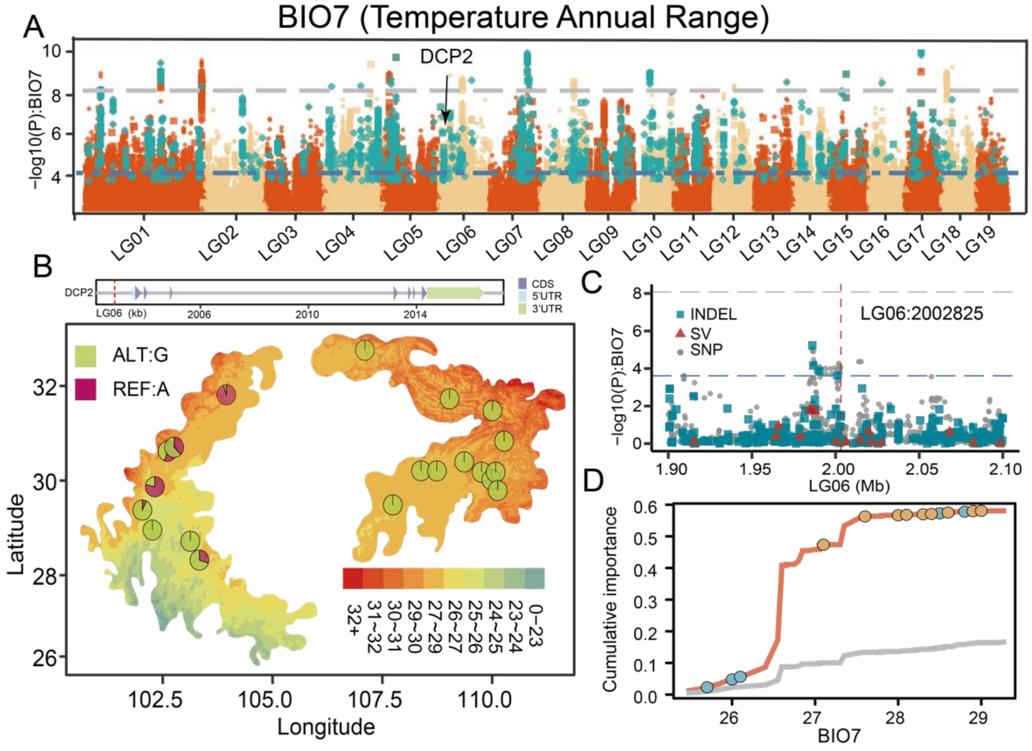

**Continuation Supplemental Figure 11.**

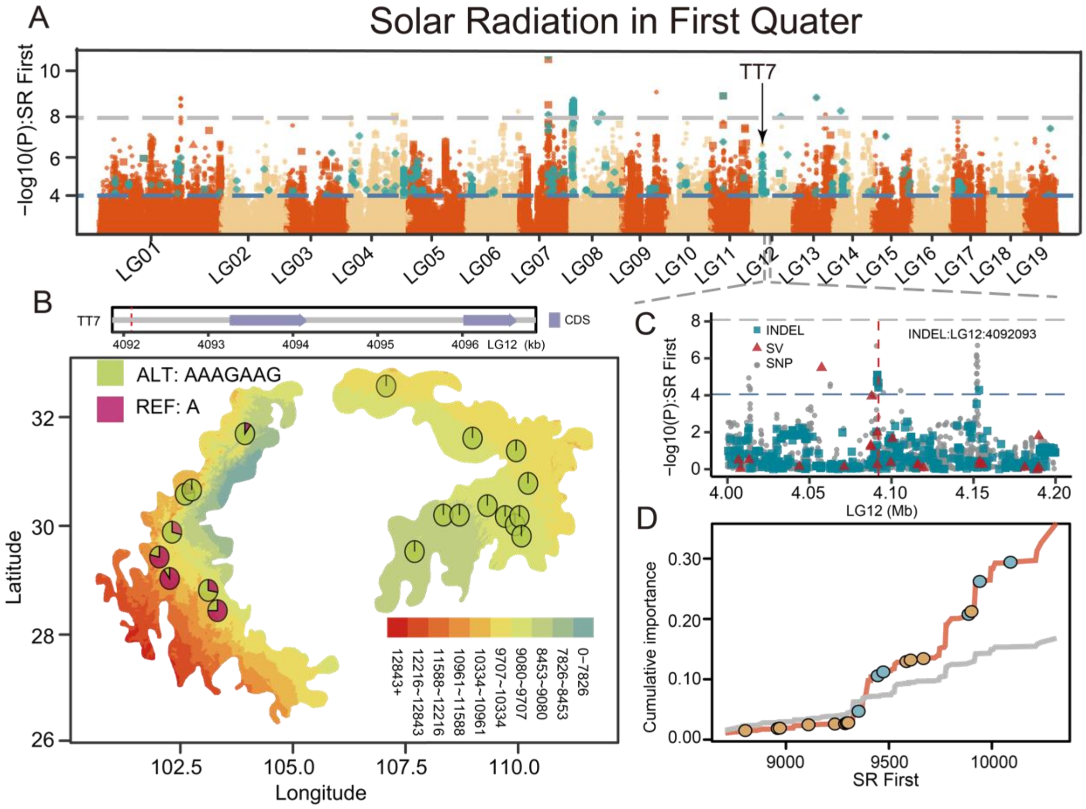

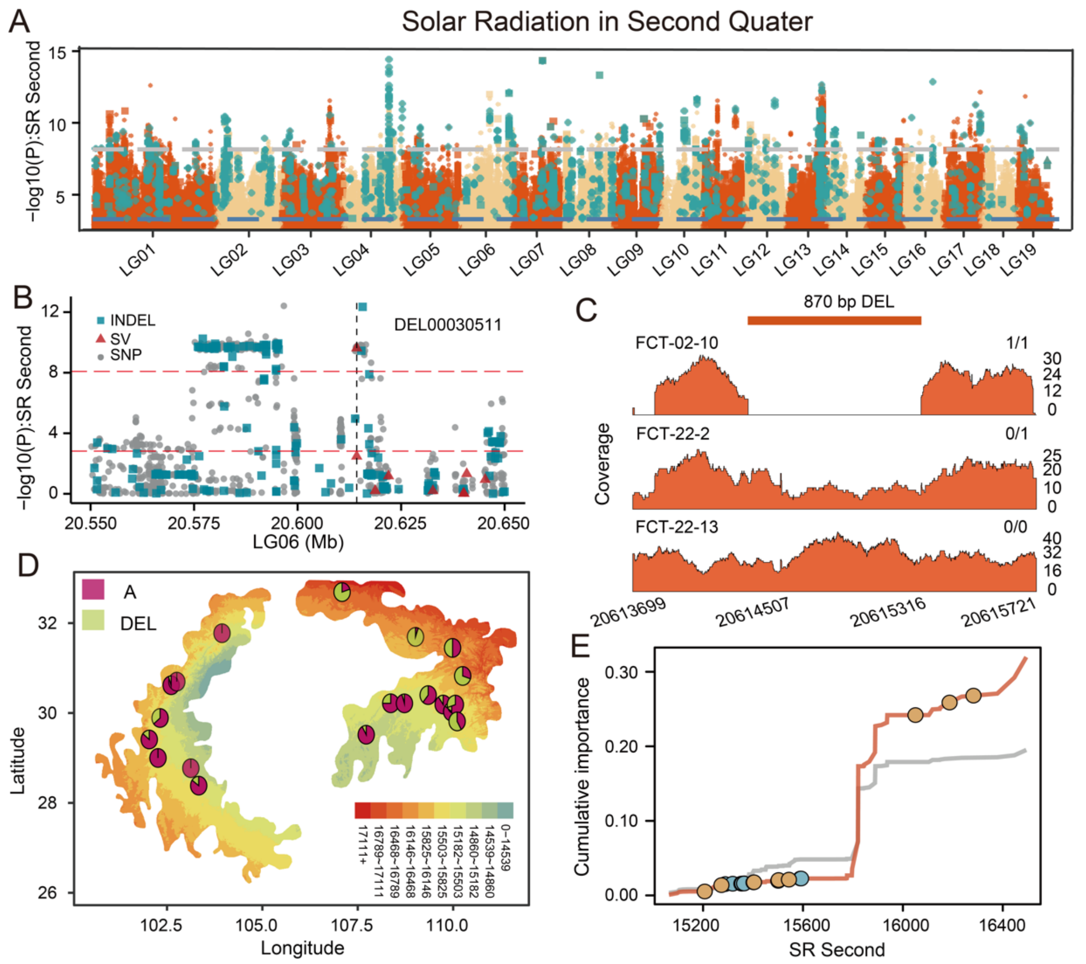


**Supplemental Figure 12.** Example of an adaptive structural variants. **(A)** Manhattan plot displaying variants associated with Solar Radiation in the Second Quarter (SR second). Dashed horizontal lines indicate significance thresholds (blue represents the FDR correction 0.05; grey line represents the Bonferroni correction, adjusted *P value* =0.05). The selected candidate gene is labeled at the corresponding genomic position. **(B)** Local magnification of the Manhattan plots around the selected genes. **(C)** A putative deletion variant, 870bp in length, associated with SR second, visualized using Samplot with Illumina short-read sequencing evidence**.** **(D)** Allele frequencies of the SV across 20 populations. The map colors represent climate gradients across the distribution range under the current scenario. **(E)** Cumulative importance of allelic change along environmental gradients for neutral variants (grey line) and adaptive variants associated with SR second (orange line).

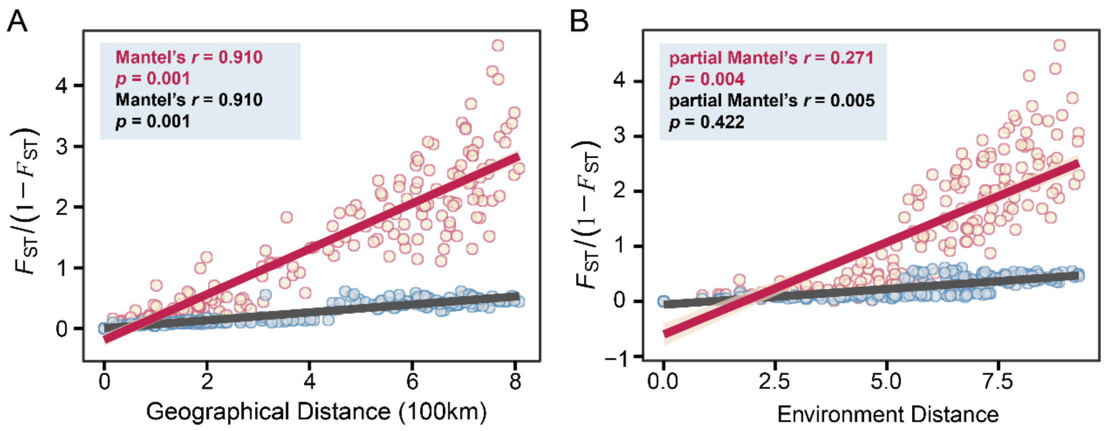


**Supplemental Figure 13. (A)** IBD analyses (Mantel test, two-sided) of pairwise populations based on neutral (blue dots and grey line) and adaptive variants (red dots and red line) separately. **(B)** IBE analyses (partial Mantel test, two-sided, controlling for the effect of geographic distance) of pairwise populations based on neutral (blue dots and grey line) and adaptive variants (red dots and red line) separately. The orange and grey shadow of linear regressions denote the 95% confidence intervals.

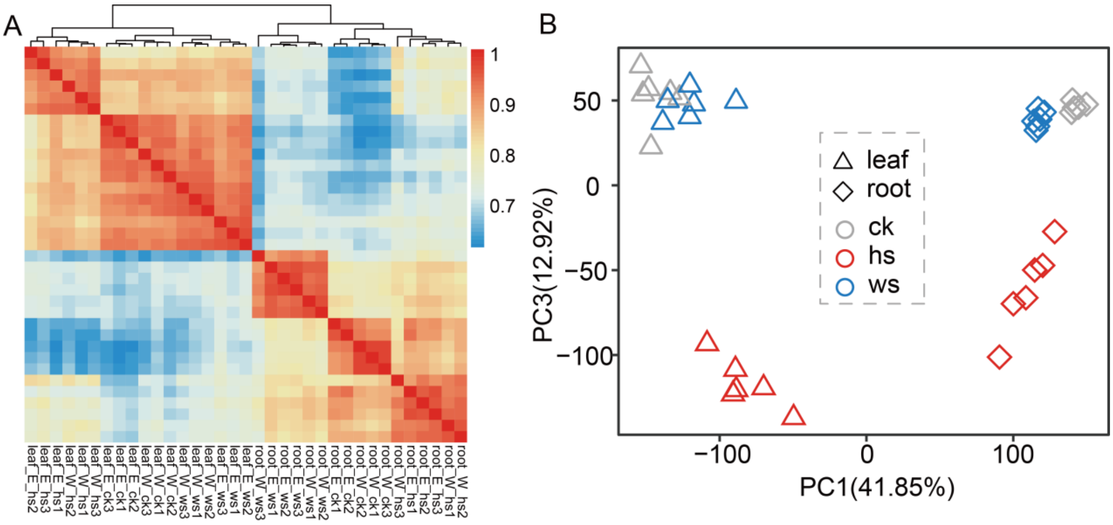


**Supplemental Figure 14.** Gene expression patterns of *P.lasiocarpa* based on hierarchical clustering **(A)** and PCA **(B)** across samples from different tissues and under various stress conditions.

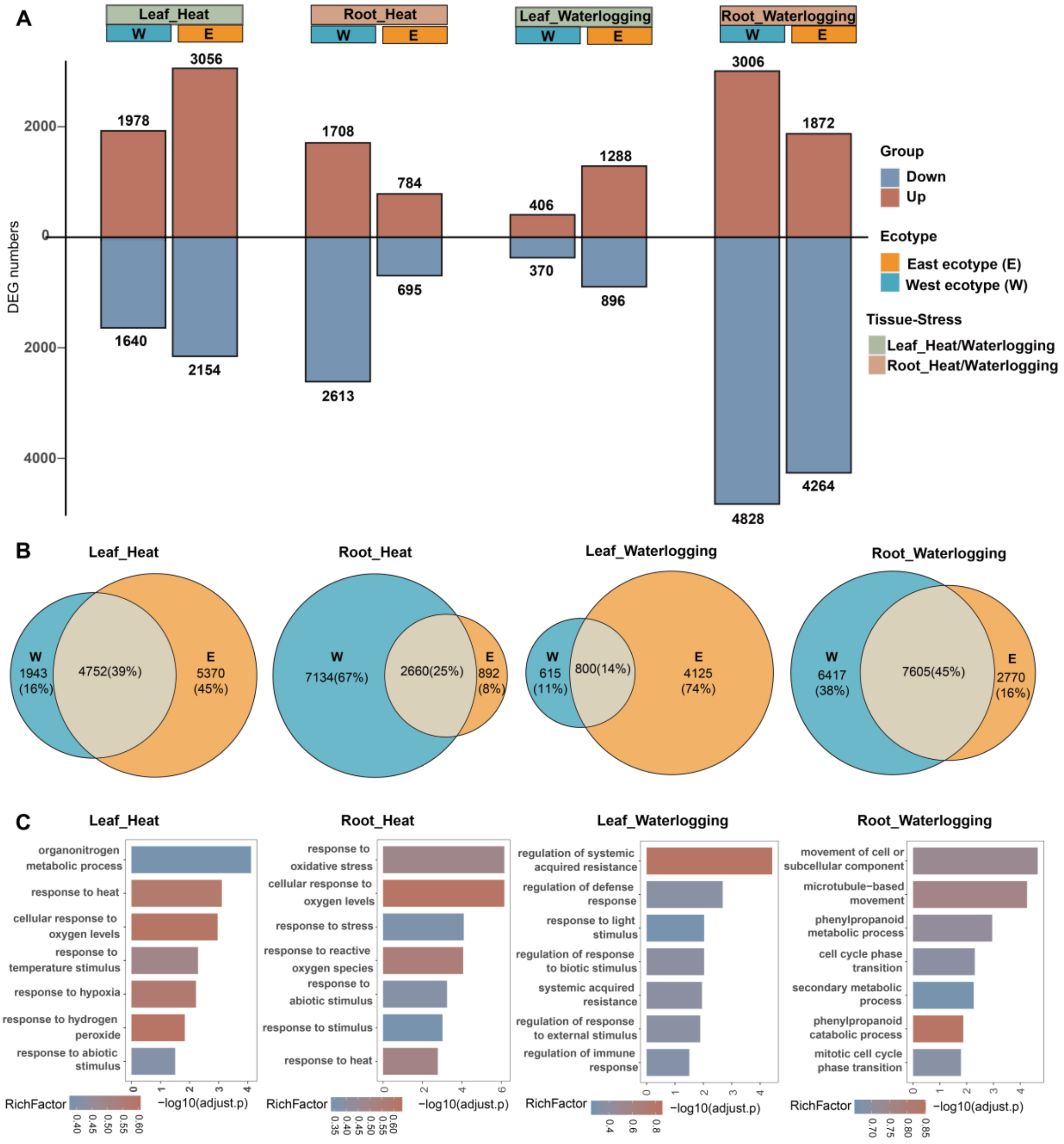


**Supplemental Figure 15.** The differential expression profiles under heat stress and submerge stress in leaf and root tissues. **(A)** The bar graphs illustrate the number of up- and down-regulated genes. The total number of differentially expressed genes (DEGs) under different condition is provided at the top of each bar. **(B)** Venn diagram showing the number of unique and shared differentially expressed genes among leaf and root tissues. (C) GO term enrichment analysis of DEGs. Significantly enriched GO terms are shown as histograms.

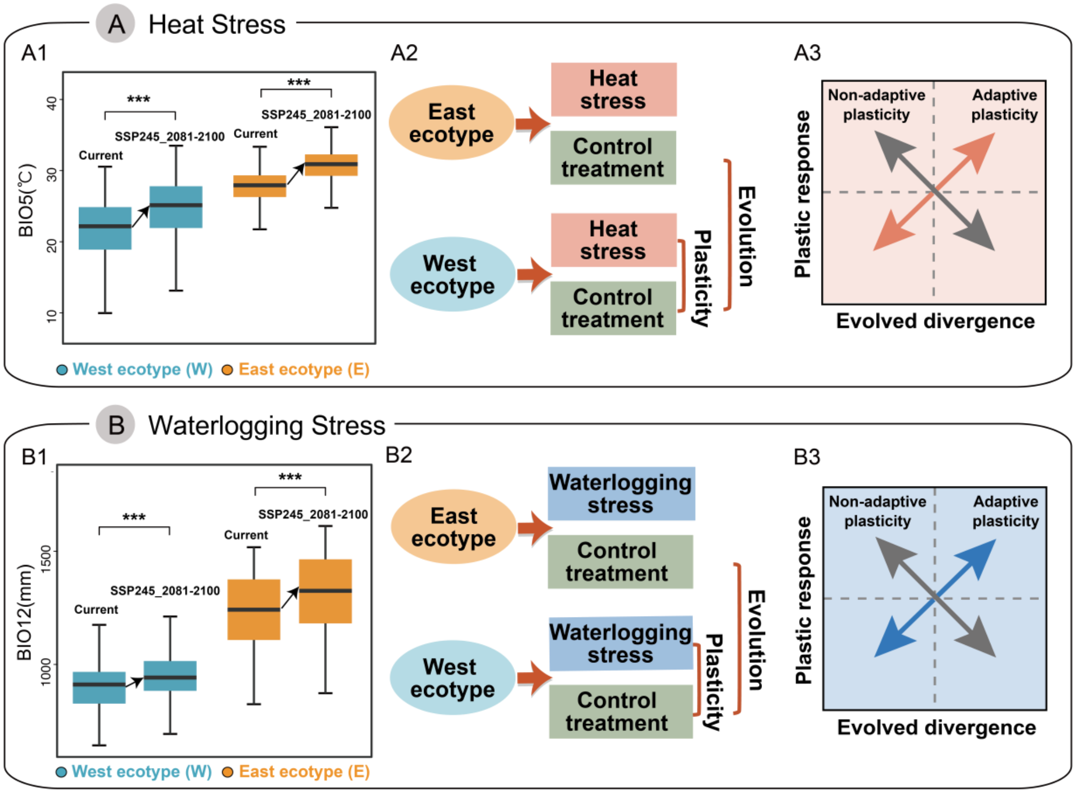


**Supplemental Figure 16.** Conceptual figure depicting the gene expression plasticity and evolved expression divergence. East and West ecotype exhibit significant differences in the environmental variables: Maximum Temperature of Warmest Month (BIO5, **A1**) and Annual Precipitation (BIO12, **B1**) in current and future period (SSP245_2081-2100). The framework shows the experimental design for heat stress **(A2)** and submerge stress **(B2)**. Plasticity is elucidated by comparing control treatments with abiotic stress treatments, while evolution is illustrated by comparing control treatment plants between the two ecotypes. In experiments involving heat stress **(A3)** and submerge stress **(B3)**, adaptive plasticity is indicated when evolved changes in gene expression (*x*-axis) positively correlate with plastic changes in gene expression (*y*-axis), and negatively correlated plasticity is non-adaptive.

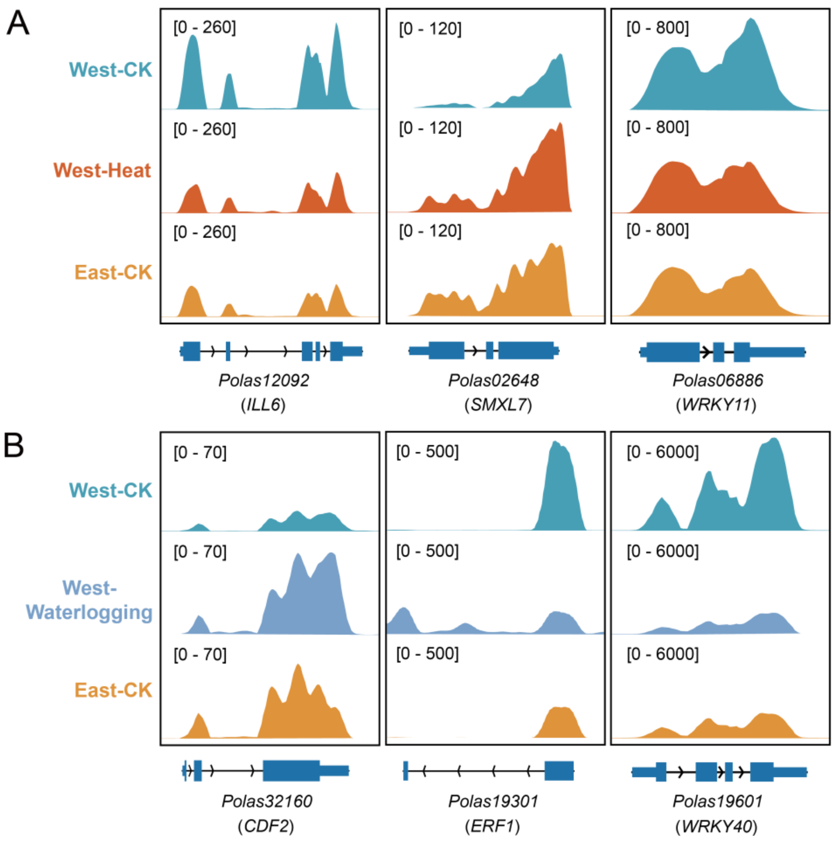


**Supplemental Figure 17.** Genomic tracks showing the expression of three representative genes that display adaptive gene expression plasticity in response to heat **(A)** and submerge stresses **(B)**.

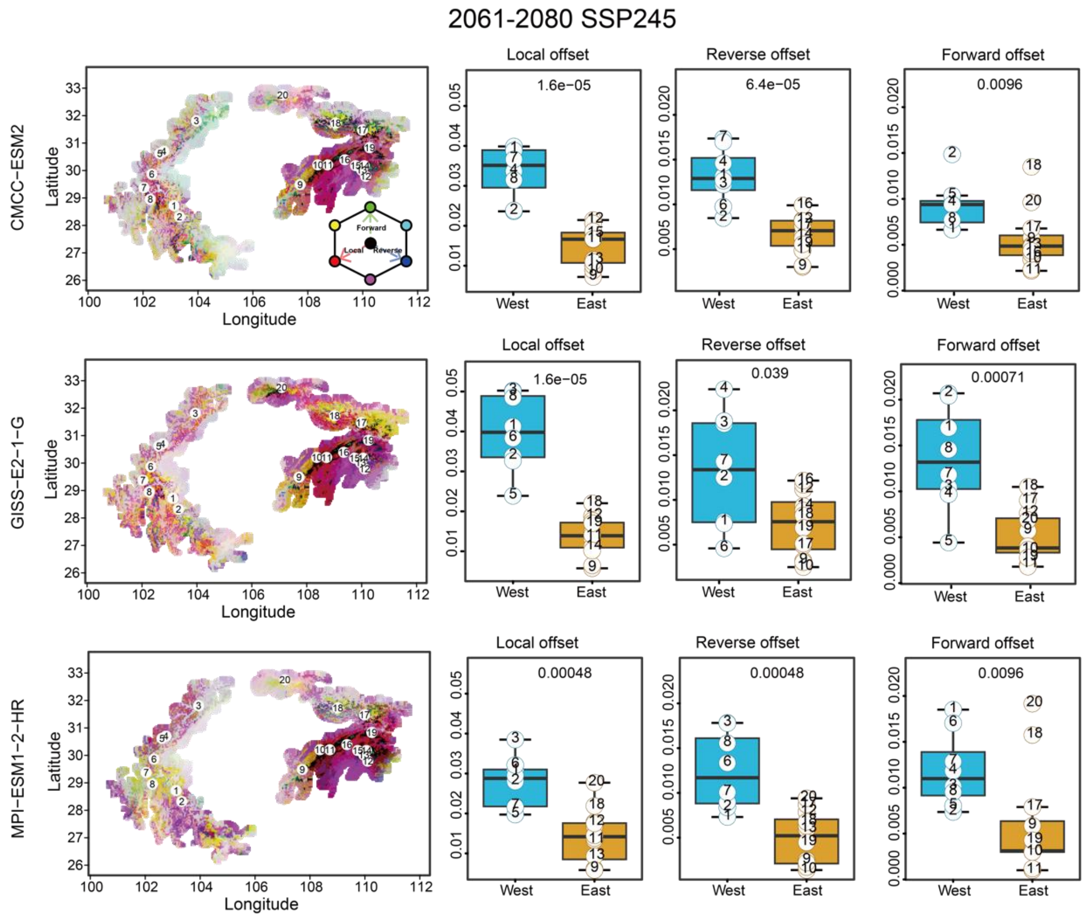


**Supplemental Figure 18**. Comparison of genetic offset estimates (local offset: red; forward offset: green; and reverse offset: blue) based on 15 BIO variables with adaptive variants identified in the natural distribution of *P. lasiocarpa* using three different climate models (CMCC-ESM2, GISS-E2-1-G, MPI-ESM1-2-HR) under moderate (SSP245) and severe (SSP585) shared socioeconomic pathways for the years 2080 (2061-2080) and 2100 (2081-2100). Circles on the map represent sampled populations. The boxplots compare local, reverse, and forward offsets between western and eastern populations, with the *P-value* displayed at the top of the boxplots.

**Continuation supplemental Figure 18.**

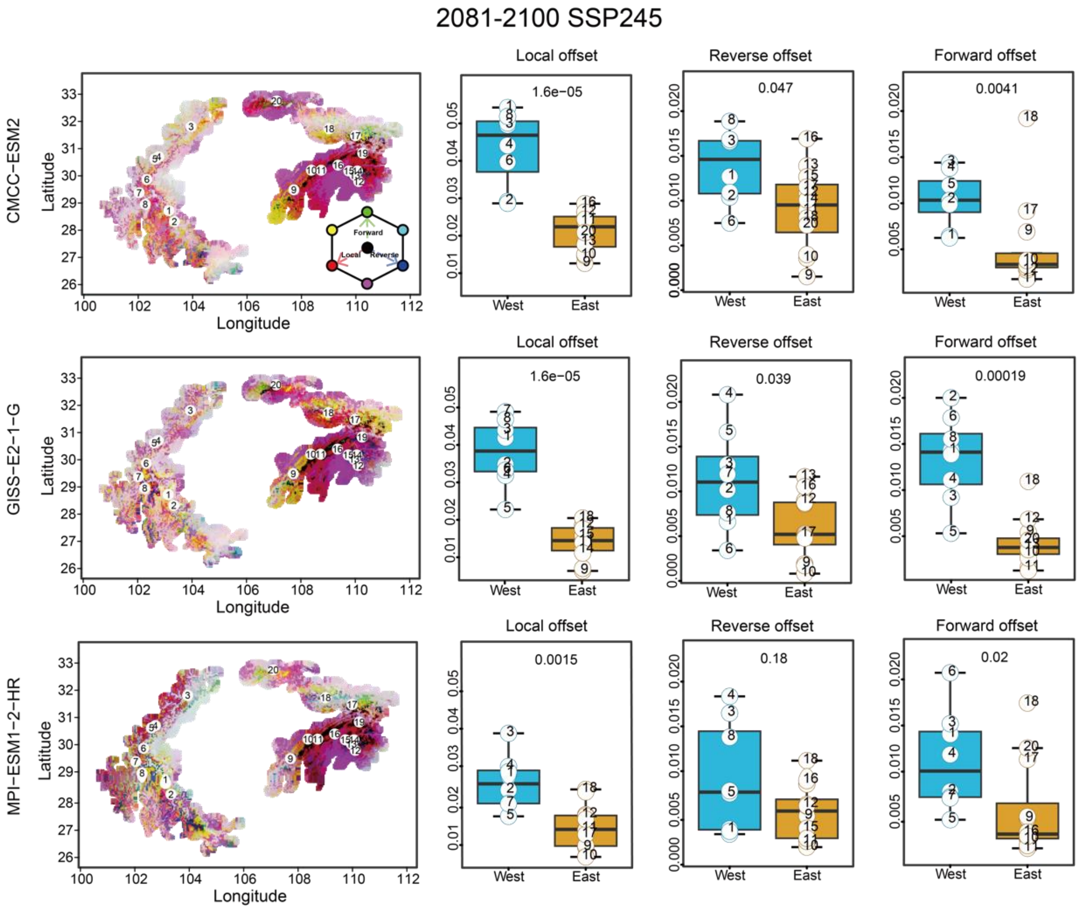

**Continuation supplemental Figure 18.**

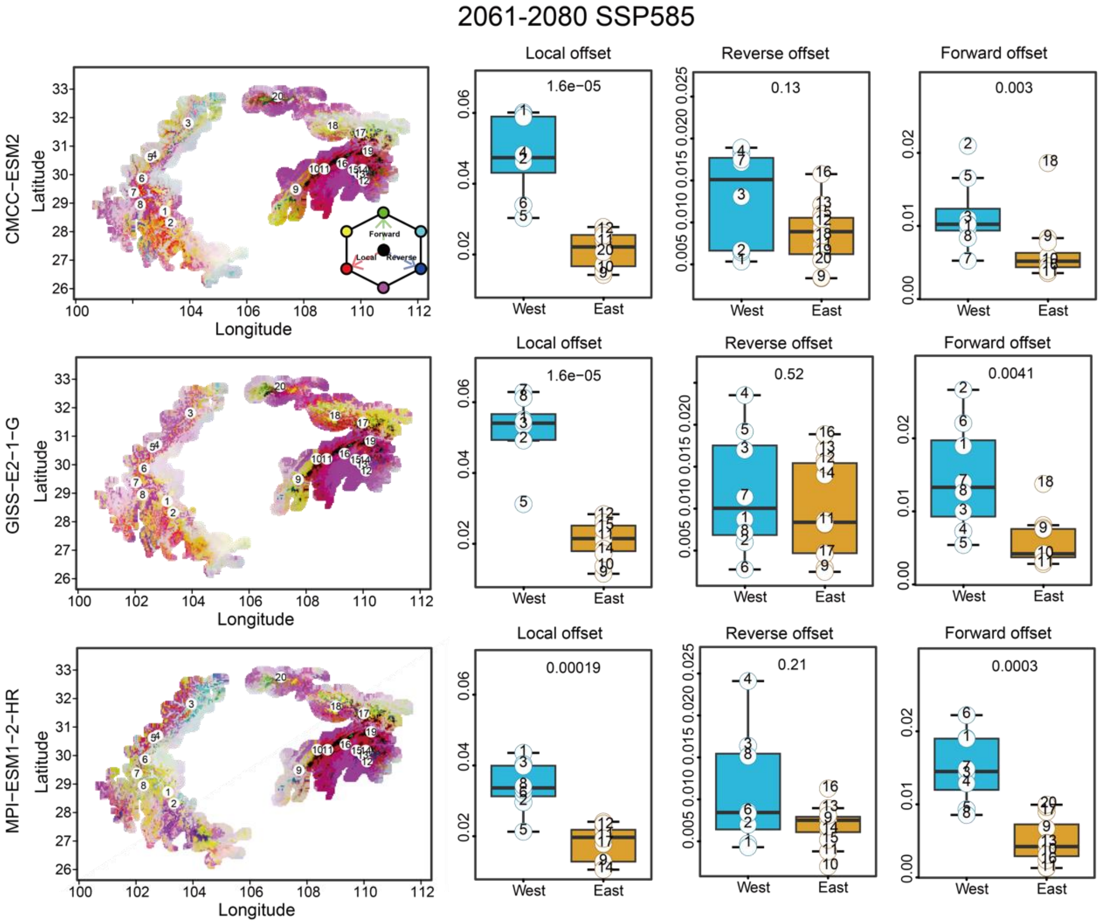

**Continuation supplemental Figure 18.**

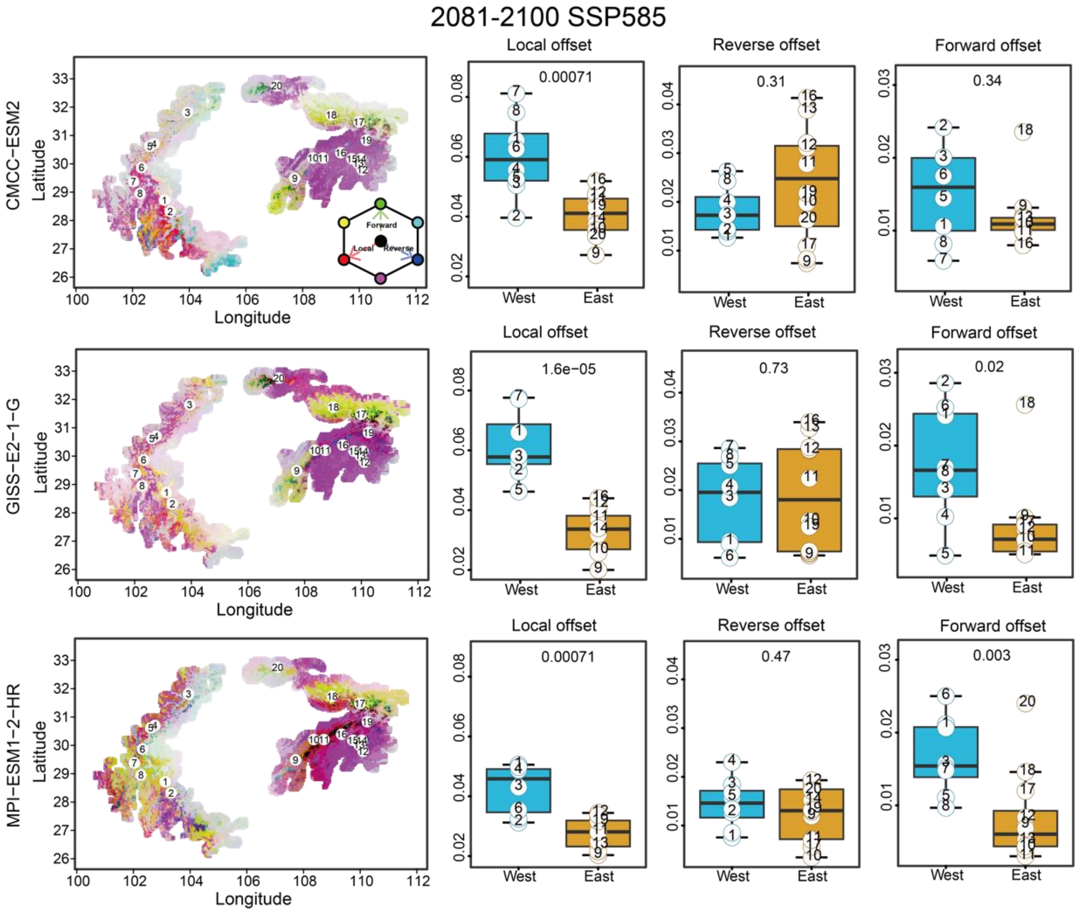

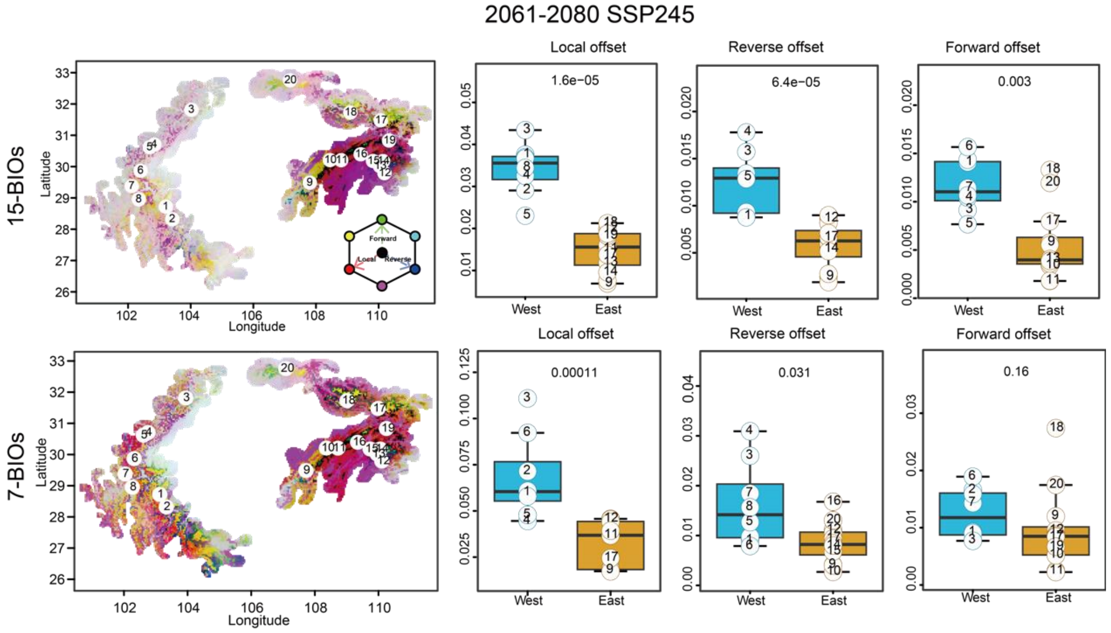


**Supplemental Figure 19.** Comparison of average genetic offsets across three climate models using 15 BIO variables and 7 uncorrelated BIO variables (i.e. BIO1, BIO4, BIO7, BIO8, BIO10, BIO15 and BIO18) under moderate (SSP245) and severe (SSP585) shared socioeconomic pathways for the years 2080 (2061-2080) and 2100 (2081-2100). Circles on the map represent sampled populations. Boxplots compare local, reverse, and forward offsets between western and eastern populations, with the *P*-value displayed at the top of the boxplots.

**Continuation supplemental Figure 19.**

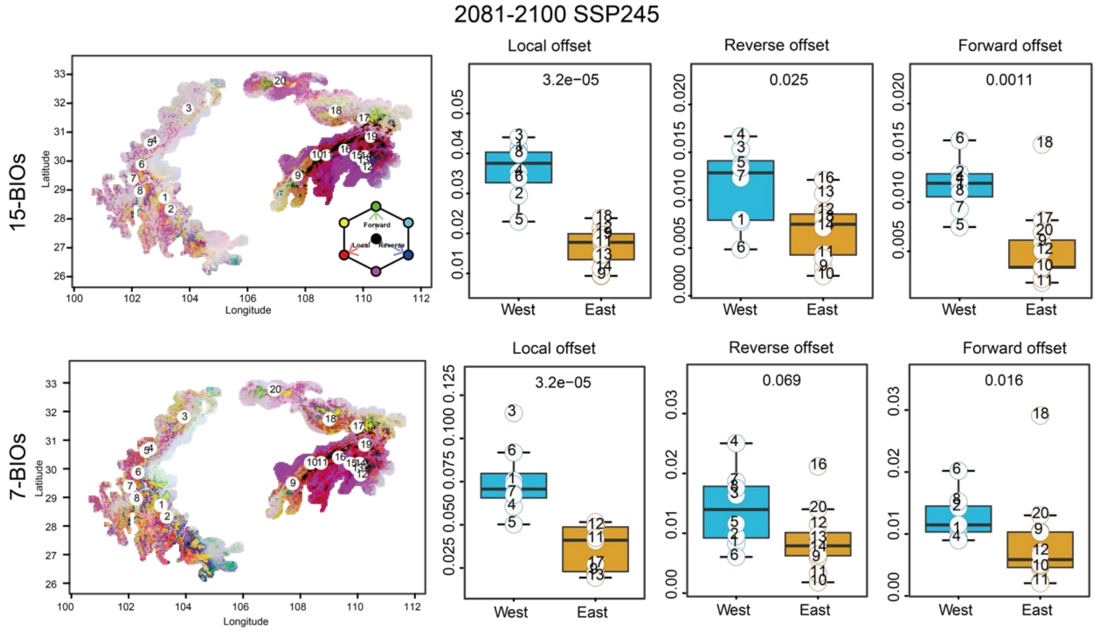

**Continuation supplemental Figure 19.**

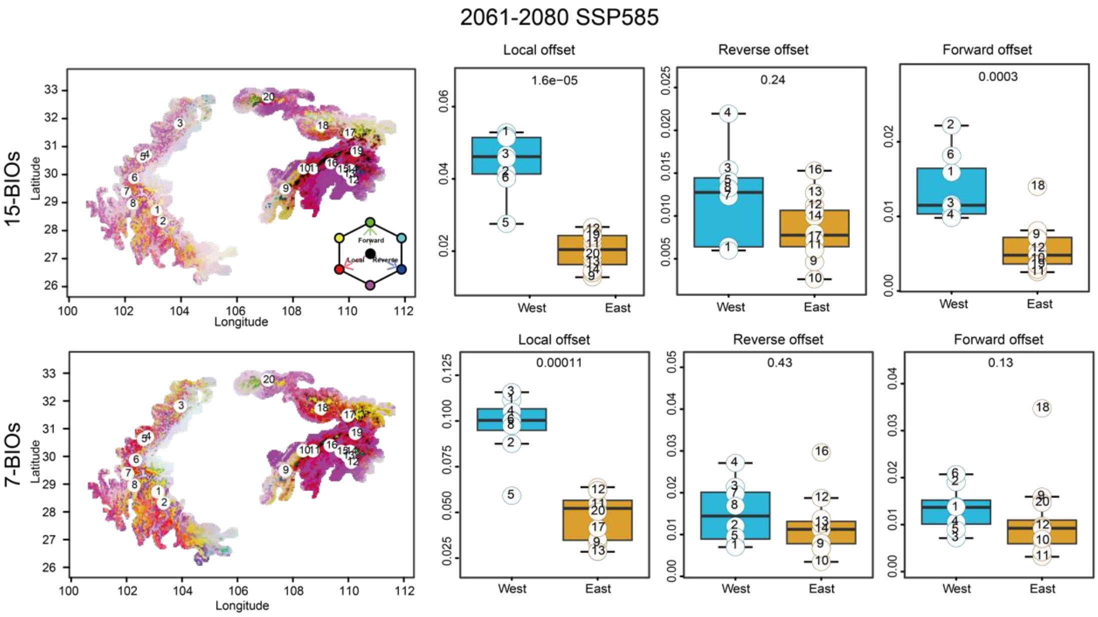

**Continuation supplemental Figure 19.**

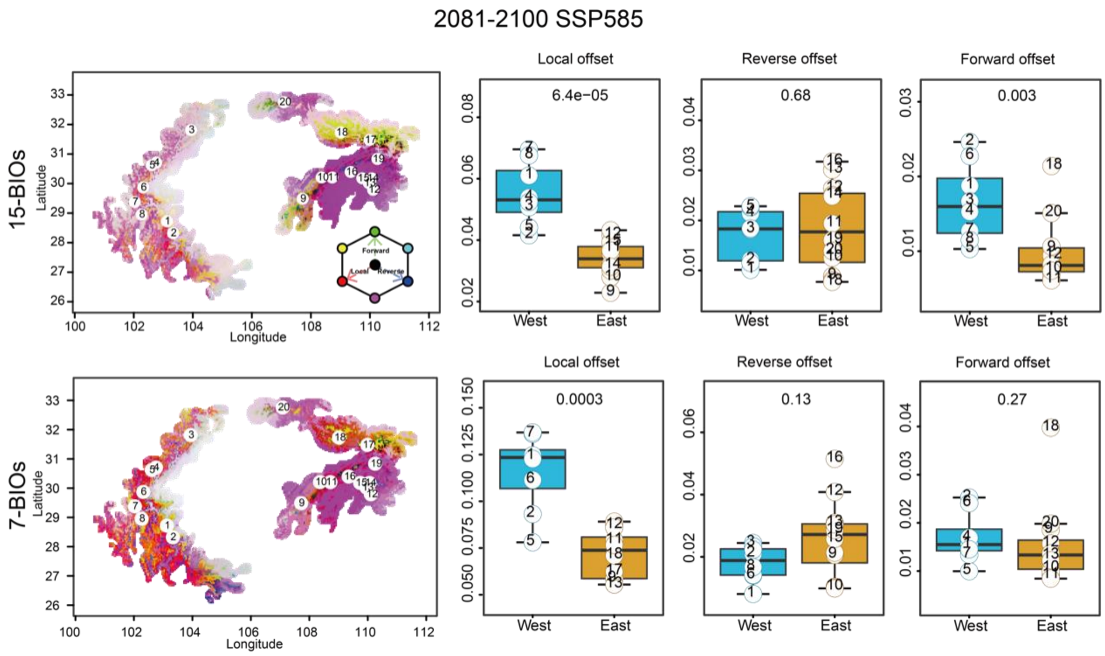

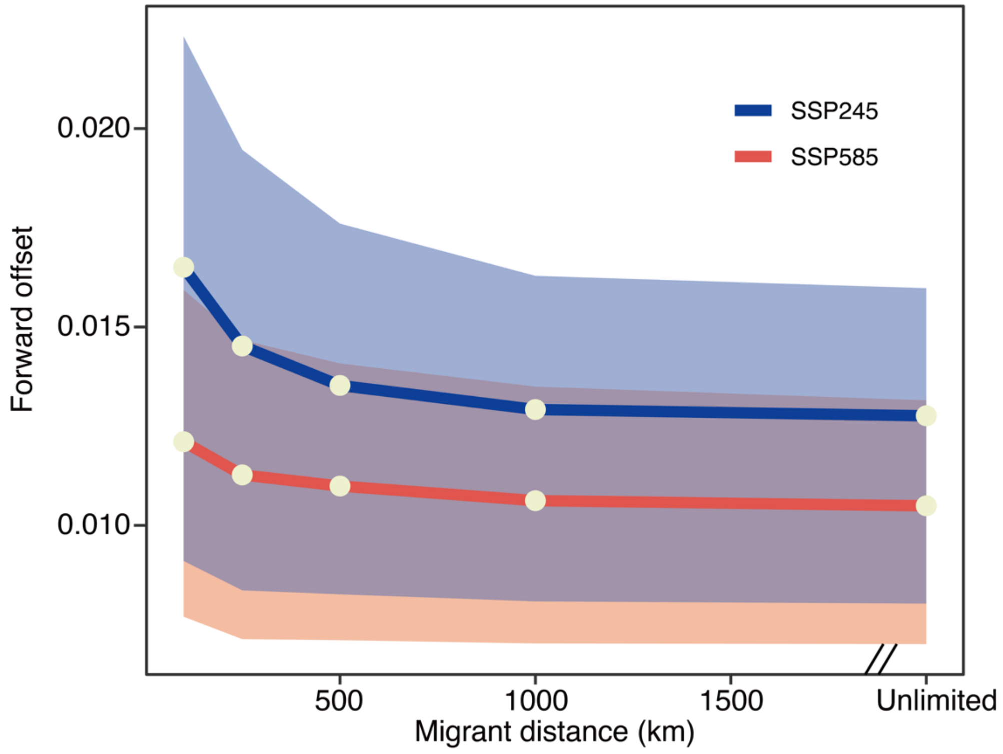


**Supplemental Figure 20.** Effect of maximum dispersal distance on the estimates of forward offset. The forward offset is shown for two SSPs for the period 2061-2080. The bands represent the range between the 25th and 75th percentiles, with the points indicating the median values.

**Supplementary Table 1 Statistics of sequencing reads and genome survey of *Populus lasiocarpa* genome.**

| **Platform** | **Total reads** | **Total bases** | **Coverage (x)** |
|---|---|---|---|
| Nanopore | 2,477,672 | 49,951,661,793 | 119.06 |
| Illumina | 242,486,812 | 36,373,021,800 | 86.70 |
| Hi-C | 416,861,438 | 62,529,215,700 | 149.04 |
| Kmer | | 17 | |
| Estimated genome size | | 451.20 Mb | |
| Estimated heterozygous ratio | | 0.6% | |

**Supplementary Table 2. Statistics of assembled contigs for *P. lasiocarpa* genome.**

| | **Preliminary Assembly** | | **Polished Genome** | | **Nonredundant and Noncontaminated Genome** | |
|---|---|---|---|---|---|---|
| | **Length** | **Number** | **Length** | **Number** | **Length** | **Number** |
| Total | 463,889,113 | 258 | 419,540,624 | 105 | 419,540,624 | 105 |
| Max | 31,312,713 | - | 31,312,713 | - | 31,312,713 | - |
| N50 | 8,459,212 | 17 | 9,191,839 | 15 | 8,459,212 | 15 |
| N60 | 6,067,086 | 24 | 7,226,573 | 20 | 6,067,086 | 20 |
| N70 | 3,878,110 | 34 | 4,729,585 | 27 | 3,878,110 | 27 |
| N80 | 1,970,620 | 51 | 3,350,016 | 37 | 1,970,620 | 37 |
| N90 | 917,550 | 85 | 1,708,288 | 54 | 917,550 | 54 |

**Supplementary Table 3. Scaffolding of contigs based on Hi-C data.**

| Peudo-chromosome | Length（Contig number） |
|---|---|
| LG01 | 52,992,279 (9) |
| LG02 | 28,222,690 (3) |
| LG03 | 26,620,501 (7) |
| LG04 | 26,318,386 (3) |
| LG05 | 25,382,582 (6) |
| LG06 | 22,749,246 (4) |
| LG07 | 22,695,253 (7) |
| LG08 | 21,045,204 (3) |
| LG09 | 20,558,685 (7) |
| LG10 | 18,743,377 (4) |
| LG11 | 18,014,361 (3) |
| LG12 | 17,702,534 (7) |
| LG13 | 17,409,083 (5) |
| LG14 | 17,212,443 (8) |
| LG15 | 17,022,449 (4) |
| LG16 | 16,768,192 (8) |
| LG17 | 16,667,112 (2) |
| LG18 | 15,551,821 (8) |
| LG19 | 14,625,206 (1) |
| Total | 416,301,404 (99) |

**Supplementary Table 4. Statistics of BUSCO evaluation for genome assembly (Nonredundant and Noncontaminated Genome).**

| BUSCO | Number | Percentage |
|---|---|---|
| Complete BUSCOs (C) | 1,346 | 97.89% |
| Complete and single-copy BUSCOs(S) | 1,089 | 79.20% |
| Complete and duplicated BUSCOs(D) | 257 | 18.69% |
| Fragmented BUSCOs(F) | 8 | 0.58% |
| Missing BUSCOs(M) | 21 | 1.53% |
| Total BUSCO groups searched | 1,375 | 100.00% |

**Supplementary Table 5. Assessment of genome sequence consistency by Illumina short-read mapping rate.**

| | | Percentage (%) |
|---|---|---|
| Reads | Mapping rate | 97.83 |
| | Properly mapping rate | 91.67 |
| Genome | Unique mapping coverage | 98.55 |
| | Coverage at least 5x | 98.08 |
| | Coverage at least 10x | 97.51 |
| | Coverage at least 20x | 95.35 |
| | single-base accuracy (Depth>=5x) | 99.99% |

**Supplementary Table 6. Statistics of transposable elements (TEs) annotation for *P. lasiocarpa* genome.**

| | **Total size (bp)** | **Percentage of genome assembly (%)** |
|---|---|---|
| **Class I: Retrotransposon** | 87,353,143 | 20.82 |
| **LTR Retrotransposon** | 63,727,493 | 15.19 |
| Copia | 16,263,189 | 3.88 |
| Gypsy | 47,464,304 | 11.31 |
| **Non-LTR Retrotransposon** | 1,284,203 | 00.31 |
| LINE | 1,184,497 | 0.28 |
| **Class II: DNA transposon** | 66,604,631 | 15.88 |
| Helitron | 37,895,370 | 9.03 |
| CACTA | 7,305,033 | 1.74 |
| Mutator | 13,665,986 | 3.26 |
| PIF_Harbinger | 2,166,554 | 0.52 |
| Tcl_Mariner | 393,947 | 0.09 |
| hAT | 5,177,741 | 1.23 |
| **Unclassified** | 14,723,521 | 3.51 |
| **Total** | 168,681,295 | 40.2 |

**Supplementary Table 7. Statistics of the annotation of protein-coding genes for *P. lasiocarpa* genome.**

| **Gene annotation features** | |
|---|---|
| Gene number | 39,008 |
| Average gene length (bp) | 3558.82 |
| Mean exons number per mRNA | 4.86 |
| Mean CDS number per mRNA | 4.74 |
| Average CDS length (bp) | 1093.61 |
| Average intron length (bp) | 1930.11 |

**Supplementary Table 8. Evaluation of assembly completeness with respect to gene space using BUSCO.**

| Description | Number | Percentage |
|---|---|---|
| Complete BUSCOs (C) | 1,557 | 96.47% |
| Complete and single-copy BUSCOs (S) | 1,300 | 80.55% |
| Complete and duplicated BUSCOs (D) | 257 | 15.92% |
| Fragmented BUSCOs (F) | 20 | 1.24% |
| Missing BUSCOs (M) | 37 | 2.29% |
| Total BUSCO groups searched | 1,614 | 100.00% |

**Supplementary Table 9. Statistics of functional annotation of protein-coding genes of *P. lasiocarpa* genome.**

| Database | Number | Percentage (%) |
|---|---|---|
| Pfam | 26,176 | 67.10 |
| Interproscan | 33,957 | 87.05 |
| KEGG | 11,219 | 28.76 |
| NR | 35,348 | 90.62 |
| Swiss-Prot | 27,161 | 69.63 |
| KOG | 31,070 | 79.65 |
| COG | 12,590 | 32.28 |
| Tremble | 35,855 | 91.92 |
| GO | 27,698 | 71.01 |
| Unannotated | 2,355 | 6.04 |
| Total | 39008 | |

**Supplementary Table 10. Statistics of Non-coding RNA annotation of *P. lasiocarpa* genome.**

| **Types** | | **Number** | **Percentage (%)** |
|---|---|---|---|
| miRNA | | 3215 | 60.41 |
| tRNA | pesdo | 709 | 13.32 |
| | complete | 646 | 12.14 |
| rRNA | 18s_rRNA | 19 | 0.36 |
| | 28s_rRNA | 15 | 0.28 |
| | 8s_rRNA | 65 | 1.22 |
| snRNA | CD-box | 312 | 5.86 |
| | HACA-box | 81 | 1.52 |
| | Sm-class | 84 | 1.58 |
| | Lsm-class | 21 | 0.39 |
| | other | 155 | 2.91 |
| Total | | 5322 | |

**Supplementary Table 11 (Excel).** Geographical sampling information and summary statistics of whole-genome re-sequencing data for 200 samples.

**Supplementary Table 12 (Excel).** Candidate genes in balancing selection regions.

**Supplementary Table 13 (Excel).** Candidate genes in divergent selection regions.

**Supplementary Table 14.** Environmental variables used in this study and the number of environment-associated genetic variants detected by LFMM and GEMMA.

| Code | Variable | LFMM & GEMMA |
|---|---|---|
| **Temperature** | | |
| BIO1 | Annual Mean Temperature | 27 |
| BIO2 | Mean Diurnal Range | 106 |
| BIO3 | Isothermality (BIO2/BIO7) | 2 |
| BIO4 | Temperature Seasonality | 3784 |
| BIO5 | Maximum Temperature of Warmest Month | 38 |
| BIO6 | Minimum Temperature of Coldest Month | 19 |
| BIO7 | Temperature Annual Range (BIO5-BIO6) | 2359 |
| BIO8 | Mean Temperature of Wettest Quarter | 34 |
| BIO9 | Mean Temperature of Driest Quarter | 0 |
| BIO10 | Mean Temperature of Warmest Quarter | 116 |
| BIO11 | Mean Temperature of Coldest Quarter | 0 |
| **Precipitation** | | |
| BIO12 | Annual Precipitation | 5953 |
| BIO13 | Precipitation of Wettest Month | 0 |
| BIO14 | Precipitation of Driest Month | 7310 |
| BIO15 | Precipitation Seasonality | 353 |
| BIO16 | Precipitation of Wettest Quarter | 88 |
| BIO17 | Precipitation of Driest Quarter | 9691 |
| BIO18 | Precipitation of Warmest Quarter | 101 |
| BIO19 | Precipitation of Coldest Quarter | 9694 |
| **Other** | | |
| CEC | Cation exchange capapcity(at ph 7) of soil | 250 |
| OCD | Organic carbon density of soil | 0 |
| Nitrogen | Nitrogen of soil | 0 |
| Vapr Annual | water vapor pressure (kPa) | 396 |
| PH | pH of soil in H2O | 0 |
| SR Annual | Mean annual UV radiation | 0 |
| SR First | Mean UV radiation in the first quarter | 511 |
| SR Second | Mean UV radiation in the second quarter | 4978 |
| SR Third | Mean UV radiation in the third quarter | 0 |
| SR Fourth | Mean UV radiation in the fourth quarter | 0 |

**Supplementary Table 15.** Significant gene ontology (GO) terms of genes associated with environmentally adaptive variants.

| GO.ID | Term | Rich factor | Fisher.p |
|---|---|---|---|
| **GO:0006559** | L-phenylalanine catabolic process | 0.667 | 0.0002 |
| **GO:1902222** | erythrose 4-phosphate/phosphoenolpyruvate family amino acid catabolic process | 0.667 | 0.0002 |
| **GO:0019439** | aromatic compound catabolic process | 0.126 | 0.0013 |
| **GO:0009074** | aromatic amino acid family catabolic process | 0.357 | 0.0014 |
| **GO:0042594** | response to starvation | 0.148 | 0.0015 |
| **GO:0006558** | L-phenylalanine metabolic process | 0.333 | 0.0019 |
| **GO:1902221** | erythrose 4-phosphate/phosphoenolpyruvate family amino acid metabolic process | 0.333 | 0.0019 |
| **GO:0022900** | electron transport chain | 0.172 | 0.0024 |
| **GO:0000165** | MAPK cascade | 0.600 | 0.0024 |
| **GO:0009772** | photosynthetic electron transport in photosystem II | 0.600 | 0.0024 |
| **GO:1901361** | organic cyclic compound catabolic process | 0.119 | 0.0026 |
| **GO:0031667** | response to nutrient levels | 0.138 | 0.0031 |
| **GO:1905613** | regulation of developmental vegetative growth | 1.000 | 0.0042 |
| **GO:1905615** | positive regulation of developmental vegetative growth | 1.000 | 0.0042 |
| **GO:0046274** | lignin catabolic process | 0.231 | 0.0053 |
| **GO:0046271** | phenylpropanoid catabolic process | 0.222 | 0.0065 |
| **GO:0009991** | response to extracellular stimulus | 0.125 | 0.0065 |
| **GO:0009267** | cellular response to starvation | 0.149 | 0.0075 |
| **GO:0080186** | developmental vegetative growth | 0.429 | 0.0077 |
| **GO:0006638** | neutral lipid metabolic process | 0.238 | 0.0094 |
| **GO:0006639** | acylglycerol metabolic process | 0.238 | 0.0094 |
| **GO:0033045** | regulation of sister chromatid segregation | 0.238 | 0.0094 |

**Supplementary Table 16.** Significant gene ontology (GO) terms of genes in haploblock regions.

| GO.ID | Term | Rich factor | Fisher.p |
|---|---|---|---|
| GO:0030307 | positive regulation of cell growth | 0.800 | 1.70E-07 |
| GO:0010105 | negative regulation of ethylene-activated signaling pathway | 0.286 | 3.00E-05 |
| GO:0070298 | negative regulation of phosphorelay signal transduction system | 0.286 | 3.00E-05 |
| GO:0010104 | regulation of ethylene-activated signaling pathway | 0.250 | 5.40E-05 |
| GO:0070297 | regulation of phosphorelay signal transduction system | 0.250 | 5.40E-05 |
| GO:0048440 | carpel development | 0.235 | 7.00E-05 |
| GO:0048467 | gynoecium development | 0.235 | 7.00E-05 |
| GO:1902532 | negative regulation of intracellular signal transduction | 0.200 | 0.0001 |
| GO:0015074 | DNA integration | 0.059 | 0.0003 |
| GO:0006259 | DNA metabolic process | 0.032 | 0.0005 |
| GO:0035670 | plant-type ovary development | 0.214 | 0.0008 |
| GO:0045927 | positive regulation of growth | 0.121 | 0.0010 |
| GO:0009873 | ethylene-activated signaling pathway | 0.114 | 0.0013 |
| GO:0071369 | cellular response to ethylene stimulus | 0.108 | 0.0016 |
| GO:0022900 | electron transport chain | 0.078 | 0.0018 |
| GO:0051097 | negative regulation of helicase activity | 0.400 | 0.0018 |
| GO:1905462 | regulation of DNA duplex unwinding | 0.400 | 0.0018 |
| GO:1905463 | negative regulation of DNA duplex unwinding | 0.400 | 0.0018 |
| GO:1905774 | regulation of DNA helicase activity | 0.400 | 0.0018 |
| GO:1905775 | negative regulation of DNA helicase activity | 0.400 | 0.0018 |
| GO:0001558 | regulation of cell growth | 0.103 | 0.0019 |
| GO:0010022 | meristem determinacy | 0.333 | 0.0027 |
| GO:0010582 | floral meristem determinacy | 0.333 | 0.0027 |
| GO:0051095 | regulation of helicase activity | 0.333 | 0.0027 |
| GO:0016036 | cellular response to phosphate starvation | 0.093 | 0.0027 |
| GO:1902531 | regulation of intracellular signal transduction | 0.089 | 0.0032 |
| GO:0009410 | response to xenobiotic stimulus | 0.250 | 0.0049 |
| GO:0022904 | respiratory electron transport chain | 0.115 | 0.0052 |
| GO:0010116 | positive regulation of abscisic acid biosynthetic process | 0.222 | 0.0063 |
| GO:0000160 | phosphorelay signal transduction system | 0.049 | 0.0068 |
| GO:0015969 | guanosine tetraphosphate metabolic process | 0.200 | 0.0078 |
| GO:0034035 | purine ribonucleoside bisphosphate metabolic process | 0.200 | 0.0078 |
| GO:0048438 | floral whorl development | 0.066 | 0.0096 |
